# Singletrome: A method to analyze and enhance the transcriptome with long noncoding RNAs for single cell analysis

**DOI:** 10.1101/2022.10.31.514182

**Authors:** Raza Ur Rahman, Iftikhar Ahmad, Zixiu Li, Robert Sparks, Amel Ben Saad, Alan Mullen

## Abstract

Single cell RNA sequencing (scRNA-seq) has revolutionized the study of gene expression in individual cell types, but scRNA-seq studies have focused primarily on expression of protein-coding genes. Long noncoding RNAs (lncRNAs) are more diverse than protein-coding genes, yet remain underexplored in part because they are under-represented in reference annotations applied to scRNA-seq. Merging annotations containing protein-coding and lncRNA genes is not sufficient, because the addition of lncRNA genes that overlap in sense and antisense with protein-coding genes will affect how reads are counted for both protein-coding and lncRNA genes. Here, we introduce Singletrome, a Singularity image that integrates protein-coding and lncRNA gene transfer format (GTF) annotations to generate enhanced annotations that take into account the sense and antisense overlap of annotated genes, maps scRNA-seq data, and produces files for downstream analysis and visualization. With Singletrome, we observed an increase in the number of reads mapped to exons, detected thousands of lncRNAs not included in GENCODE, and observed a decrease in uniquely mapped reads, indicating improved mapping specificity. Moreover, we were able to cluster cell types based solely on lncRNAs expression, and lncRNAs alone were able to predict cell types and human disease pathology through machine learning. This comprehensive annotation will allow mapping of lncRNA expression across cell types of the human body, facilitating the development of an atlas of human lncRNAs in health and disease with the ability to integrate new lncRNA annotations as they become available.

## Introduction

Long noncoding RNAs (lncRNAs) comprise a diverse class of transcripts that regulate pathology, including cancer (Huang et al. 2017), immunity (Kotzin et al. 2016), and liver disease (Mahpour and Mullen 2021). LncRNA transcripts are at least 200 nucleotides in length, 5’ capped, 3’ polyadenylated, and are not known to code for proteins (Statello et al. 2021). The functions of individual lncRNAs are diverse, with new activities described as additional lncRNAs are investigated. For example, *Evx1as* regulates mesendoderm differentiation through cis regulation of Even-skipped homeobox 1 (*Evx1*) (Bell et al. 2016). *DIGIT* (*GSC-DT*) interacts with BRD3 to control definitive endoderm differentiation (Daneshvar et al. 2020), and *Morrbid* controls the lifespan of immune cells by regulating the transcription of the apoptotic gene *Bcl2l11* through the enrichment of the PRC2 complex at its promoter (Kotzin et al. 2016).

Many lncRNAs also exhibit cell-type-specific patterns of expression (Liu et al. 2016). For example, *LOC646329* is enriched in single radial glia of the human neocortex (Liu et al. 2016), and *Lnc18q22.2* is induced only in hepatocytes in the setting of metabolic dysfunction-associated steatohepatitis (MASH/NASH) (Atanasovska et al. 2017). Cell-type-specific expression patterns observed for many lncRNAs suggest that lncRNA expression could support distinct clustering of cell types in single cell data.

Despite advances in our understanding of the functions of many lncRNAs and frequent examples of cell-type-specific expression, lncRNA discovery is still at a preliminary stage, and there is not yet consensus on the number of lncRNAs in the human genome. GENCODE (v32), the most widely applied genome annotation for human scRNA-seq analysis, contains 16,849 lncRNA genes (Frankish et al. 2019), but databases such as LncExpDB and Noncode now report over 100,000 human lncRNA genes (Li et al. 2021; Fang et al. 2018).

Increasing the number of lncRNAs identified in single cell data cannot be achieved by simply creating new annotations that contain known protein-coding and lncRNA genes, because the addition of tens of thousands of new genes will affect how gene expression is quantified. Current pipelines such as Cell Ranger (Zheng et al. 2017) exclude reads mapping to exons that overlap on the same strand, therefore expanding the number of annotated lncRNA exons may lead to exclusion of additional reads from an increased number of overlapping exons. Furthermore, the assignment of reads that align to antisense transcripts is challenging in part because library preparation artifacts can generate antisense reads at low frequency. For example, the widely used dUTP protocol for stranded RNA-seq (Parkhomchuk et al. 2009) can generate spurious antisense reads ranging from 0.6-3% of the sense signal (Zeng and Mortazavi 2012; Jiang et al. 2011). Analysis of 199 strand-specific RNA-seq datasets discovered that spurious antisense reads are detected in these experiments at levels greater than 1% of sense gene expression levels (Mourão et al. 2019). Additionally, mis-priming of internal poly-A tracts on RNA or template switching into the poly-T linker have been proposed as possible sources of intronic and antisense reads in single cell gene expression data (Ding et al. 2020). Ultimately, full-length RNA molecule sequencing will help to define authentic antisense RNAs. However, reverse transcriptase-based approaches are predominantly used for sequencing, and special attention needs to be directed towards distinguishing authentic antisense lncRNAs from experimental artifacts, as lncRNAs tend to be expressed at ∼10-fold lower levels than protein-coding genes (Cabili et al. 2011; Derrien et al. 2012). It is crucial to develop an approach to minimize the possibility of interpreting the presence of reads antisense to a protein-coding exon as evidence of lncRNA expression if reads are the product of library preparation.

While efforts have been made to analyze lncRNAs in scRNA-seq data for a particular set of transcripts, cell types, or datasets (Liu et al. 2016; Luo et al. 2021), no systematic efforts have been made to analyze all annotated lncRNAs in scRNA-seq data. Furthermore, these efforts do not provide a unified framework to analyze lncRNAs in scRNA-seq data. The most widely used genome annotation for scRNA-seq analysis is GENCODE, which contains only a fraction of annotated lncRNAs in the human genome. Here we develop Singletrome, a framework to create a comprehensive genome annotation of 110,599 genes consisting of 19,384 protein-coding genes from GENCODE and 91,215 lncRNA genes from LncExpDB, which takes into account the sense and antisense relationship between lncRNAs and protein-coding genes and the distribution of reads across lncRNA transcripts in each dataset. Singletrome is a Singularity image that takes two GTF annotations as input, one containing protein-coding genes and the other containing lncRNAs to generate an enhanced genome annotation. It enables browser extensible data (BED) file creation for downstream analysis, executes Cell Ranger for scRNA-seq data mapping, and merges multiple samples into a single binary alignment map (BAM) file for quality control analysis (RSeQC) and visualization (BigWig). Singletrome is invariant to genome annotation versions as it accepts two GTF files, providing flexibility across different annotation sources. While we applied it to LncExpDB, it can also be used with NONCODE and other lncRNA annotations. Additionally, although we have applied Singletrome only to human genome annotations, it can be extended to other organisms where lncRNAs and protein-coding genes are defined in GTF format. We applied Singletrome to analyze single cell data from PBMCs as well as healthy and diseased liver.

## Results

### Expanding lncRNA annotations in single cell analysis

In order to enhance the current genome annotation for lncRNAs in single cell analysis, we first evaluated how the integration of LncExpDB into GENCODE impacts the annotation. We identified 6309 protein-coding genes (42,868 exons) that overlap on the sense strand with 7531 lncRNA genes (24,357 exons) (Fig 1A & Table 1). We next evaluated lncRNA genes annotated antisense to protein-coding genes and found 10,492 protein-coding genes (47,057 exons) overlap on the antisense strand with 14,212 lncRNA genes (44,062 exons) (Fig 1A & Table 1). This situation is not unique to our new annotation, as 619 protein-coding genes (3514 exons) overlap on the sense strand with 516 lncRNA genes (2106 exons) and 3590 protein-coding genes (12,941 exons) overlap on the antisense strand with 3791 lncRNA genes (8809 exons) in GENCODE (Table 2). We removed the 7531 lncRNA genes from LncExpDB that overlap protein-coding genes on the sense strand (Fig 1B), as it is challenging to prove these lncRNA genes are not isoforms of the protein-coding genes or have coding potential. As a result, reads mapped to the protein-coding exons that overlap on the same strand with these lncRNAs are included to define Unique Molecular Identifier (UMI) counts for the protein-coding genes. To distinguish authentic antisense lncRNAs from potential artifacts, we developed a trimmed lncRNA genome annotation (TLGA) to retain all the non-overlapping lncRNA exonic regions (Fig. 1C). More specifically, we removed parts of lncRNA exons that coincided with protein-coding exons on the opposite strand with a buffer that included an additional 100 nucleotides in each direction. We kept lncRNA exons that were a minimum of 200 nucleotides in length after this trimming process. The approach to only count reads mapped to regions of lncRNAs that are not antisense to protein-coding genes reduces the risk of incorrectly calling an lncRNA as expressed based only on antisense reads that might have been generated during library preparation.

**Figure 1.**
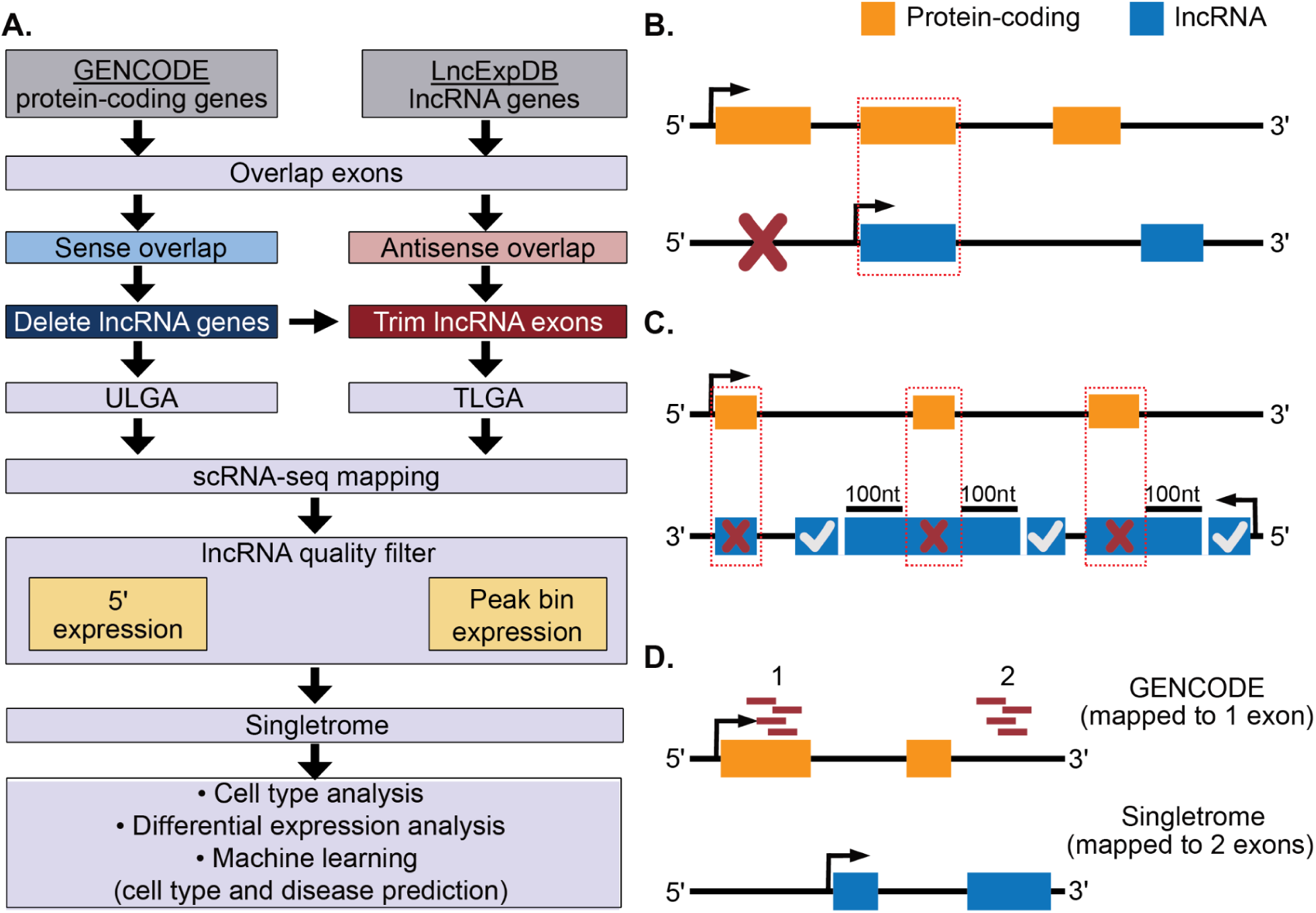
Enhancing the transcriptome with expanded lncRNA annotation for single cell analysis. **(A)** Development of Singletrome workflow. Exons of protein-coding genes from GENCODE v32 and lncRNA genes from LncExpDB v2 were integrated (Table 1). lncRNA genes overlapping on the sense strand with protein-coding genes were deleted to create the <u>u</u>ntrimmed lncRNA <u>g</u>enome <u>a</u>nnotation (ULGA), and antisense strand overlapping lncRNA exons were trimmed to create the trimmed lncRNA <u>g</u>enome <u>a</u>nnotation (TLGA) for scRNA-seq analysis. scRNA-seq data were mapped to both ULGA (to account for all lncRNA mapped reads) and TLGA (to define lncRNA expression based on reads with the highest confidence). Mapped lncRNAs were subjected to additional quality filters to remove transcripts that have reads mapped predominantly to the 5’ end of a transcript or to a single, non 3’ bin. The quality filtered lncRNAs (Singletrome) were used to perform cell type identification, differential expression analysis, and prediction of cell types and disease using machine learning. **(B)** Sense strand overlap. Cell Ranger discards reads mapped to overlapping exons on the same strand (red dotted box). To avoid miscounting reads to protein-coding genes by the inclusion of additional lncRNAs in the genome, lncRNA genes were discarded if they overlap in sense with protein-coding exons (red x), as it is more difficult to exclude the protein-coding potential of these lncRNAs. **(C)** Antisense strand overlap. Cell Ranger prioritizes alignments of sense over antisense reads. If spurious antisense reads are generated from transcripts of protein-coding genes, these could be incorrectly interpreted to indicate expression of an antisense overlapping lncRNA gene. To overcome this potential problem, we trimmed the overlapping region (red x) and an additional 100nt of lncRNA exons that were overlapping with protein-coding exons in the antisense direction. We retained the trimmed lncRNA exons if their length was at least 200nt (marked with white check). Gene and transcripts coordinates were updated accordingly. **(D)** Improved mapping specificity. Reads (red bars, 1) uniquely mapped to a single exon and were carried forward to UMI counting in GENCODE. In Singletrome, with the inclusion of 91,215 lncRNA genes (537,373 exons) these reads are now mapped to two exons (red bars 1 and 2). Removing these reads from UMI counting would improve read mapping specificity but reduce the total number of uniquely mapped reads.

**Table 1.** Integrating GENCODE v32 (Frankish et al. 2019) and LncExpDB v2 (Li et al. 2021). GENCODE (dated 27.10.2021) contains 19,384 protein-coding genes (476,299 exons), and LncExpDB (27.10.2021) contains 101,285 lncRNA genes (611,102 exons). The table indicates the number of lncRNAs that were filtered based on sense strand and antisense overlap as described in the text. * denoted no filtering of protein-coding genes and exons.

| Feature | Genes |  | Exons |  |
| --- | --- | --- | --- | --- |
| Type | protein-coding | LncRNA | protein-coding | LncRNA |
| Total | 19,384 | 101,285 | 476,299 | 611,102 |
| Sense strand overlap | 6309 | 7531 | 42,868 | 24,357 |
| Antisense strand overlap | 10,492 | 14,212 | 47,057 | 44,062 |
| Post sense strand filtering | * | 93,754 | * | 565,717 |
| Post antisense strand filtering | * | 91,215 | * | 537,373 |

**Table 2.** Distribution of protein-coding and lncRNA genes in GENCODE v32.

| Feature | Genes |  | Exons |  |
| --- | --- | --- | --- | --- |
| Type | protein-coding | LncRNA | protein-coding | LncRNA |
| Total | 19,384 | 16,849 | 476,299 | 109,075 |
| Sense strand overlap | 619 | 516 | 3514 | 2106 |
| Antisense strand overlap | 3590 | 3791 | 12,941 | 8809 |

Using this strategy, we were able to retain 11,673 of the 14,212 lncRNA genes that contain regions antisense to protein-coding genes. We deleted 2539 lncRNA genes where no exons satisfy the aforementioned criteria. Following these trimming steps we retained 91,215 of 101,285 lncRNA genes (Fig. 1C & Table 1). We then created a comprehensive genome annotation of 110,599 genes consisting of 19,384 protein-coding genes from GENCODE and 91,215 lncRNA genes containing regions that do not overlap with protein-coding genes and refer to this approach as the trimmed lncRNA <u>g</u>enome <u>a</u>nnotation (TLGA). TLGA increased the wealth of lncRNA exons by 4.93 fold (n=428,298), transcripts by 6.46 fold (n=258,106), and genes by 5.41 fold (n=74,366) compared to GENCODE. The inclusion of these additional lncRNA genes may also slightly reduce the total number of uniquely mapped reads, as some reads uniquely mapped in GENCODE will no longer be uniquely mapped with TLGA. (Fig 1D).

### Maximizing reads mapped to lncRNAs for downstream analysis

TLGA expands the number of annotated lncRNAs but still excludes regions of 11,673 lncRNAs that partially overlap antisense exons of protein-coding genes. Once we define an lncRNA as expressed in a dataset, the antisense reads could provide additional depth to assist in cell clustering and the definition of genes expressed in specific cell types for follow-up studies. In addition, lncRNAs are expressed at lower levels than protein-coding genes (Fig 2A-B and Supplementary Fig 1A-D), and antisense lncRNAs often have functional activity (Faghihi et al. 2010; Yap et al. 2010), so there are benefits to including as much information for these genes as possible once the thresholds for expression are met.

**Figure 2.**
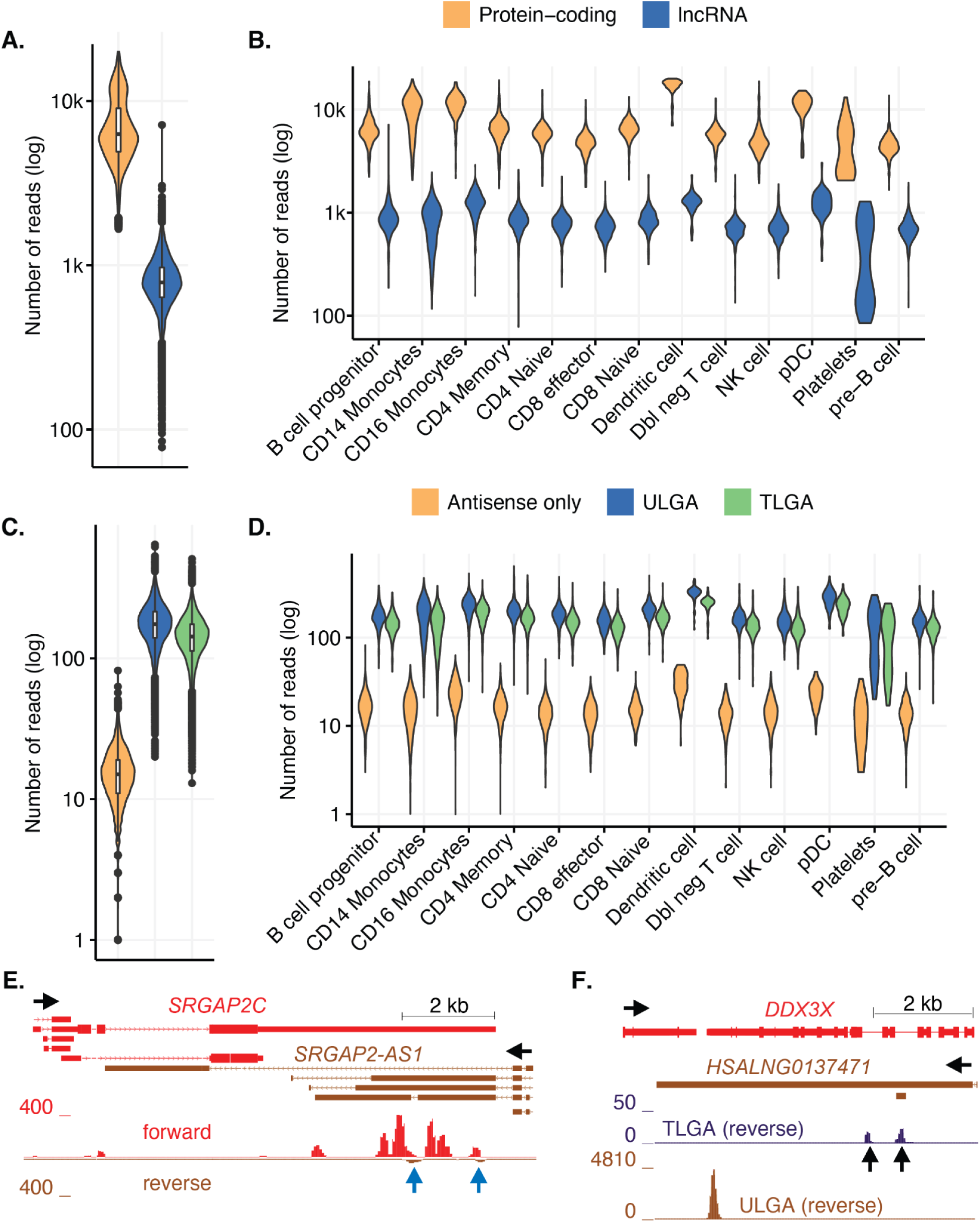
Distribution of transcripts in bulk and by cell type in PBMCs. **(A)** lncRNAs (blue) are expressed at lower levels than protein-coding genes (orange) in PBMCs. Reads aligned to lncRNAs and protein-coding genes are shown in log scale (y-axis). **(B)** lncRNAs are expressed at lower levels than protein-coding genes in all PBMC cell types. **(C)** Expression of lncRNAs that overlap protein-coding genes in the antisense direction. Expression of lncRNAs in non-overlapping regions [TLGA (green)], expression of lncRNAs in overlapping and non-overlapping regions [ULGA (blue)], and lncRNA exons that are expressed only in regions antisense to protein-coding exons [antisense only (orange)] are displayed (y-axis). The Y-axis shows the number of reads in log scale. TLGA (green) identifies lncRNAs with the highest confidence in expression but reduces the reads associated with lncRNAs as compared to ULGA (blue). **(D)** Data for each cell type are shown as described in (C). **(E)** Example where using ULGA reads could incorrectly suggest expression of an antisense lncRNA gene. *SRGAP2-AS1* (brown) is expressed antisense to *SRGAP2C* (red). Reads mapped to *SRGAP2C* (forward) are shown in red, while ULGA reads mapped to *SRGAP2-AS1* (reverse) are shown in brown. In this example, the reads mapped to *SRGAP2-AS1* are contained in exons antisense to an exon of *SRGAP2C,* where there are many more reads supporting the mRNA antisense to the lncRNA (blue arrows). This lncRNA gene is discarded because there are insufficient reads to support expression of *SRGAP2-AS1* in regions that do not overlap with exons of *SRGAP2 (SRGAP2-AS1* has no reads mapped in *TLGA)*. The genomic scale is indicated on the upper right, and the direction of transcription is indicated by horizontal black arrows. **(F)** Example where TLGA identifies reads mapped to exons of *HSALNG0137471* that do not overlap (in antisense) to exons of *DDX3X* (black vertical arrows), but does not capture the majority of reads mapped towards the 3’ end of *HSALNG0137471*. LncRNA *HSALNG0137471* (brown) is antisense to *DDX3X* (red). TLGA (reverse) only displays reads mapped to *HSALNG0137471* in exons that do not overlap (in antisense) to exons of *DDX3X*. ULGA (reverse) shows all reads mapped to *HSALNG0137471*. In this case, all reads mapped to *HSALNG0137471* [ULGA (reverse)] are used for down-stream analysis after the lncRNA is defined as expressed based on TLGA reads.

To assess the impact of trimming, we compared TLGA with an <u>u</u>ntrimmed lncRNA <u>g</u>enome <u>a</u>nnotation (ULGA). In ULGA, we deleted lncRNA genes overlapping protein-coding genes on the sense strand but included all reads for the antisense overlapping lncRNAs. We mapped PBMCs (pbmc_10k_v3 from 10x Genomics), liver set 1 (GSE115469 (MacParland et al. 2018)) and liver set 2 (GSE136103 (Ramachandran et al. 2019)) scRNA-seq data with ULGA and TLGA to assess the output from each annotation.

More than 1,000 lncRNA genes in each dataset are expressed in ULGA but have no expression in TLGA. Of 14,212 antisense overlapping lncRNAs, 1458 lncRNAs in PBMCs are expressed in ULGA but not TLGA, while 1153 and 1841 lncRNAs are expressed in ULGA but not TLGA in liver sets 1 and 2, respectively (Supplementary Table 1). Reads mapped to these lncRNA genes were aligned only to regions antisense to protein-coding exons (+100nt) in ULGA and have the potential to come only from library preparation artifacts. These lncRNAs were removed from down-stream analysis.

Of the 14,212 antisense overlapping lncRNAs, 4921 lncRNAs in PBMCs, 4194 in liver set 1, and 6675 in liver set 2 are expressed in both TLGA and ULGA. For these lncRNAs, TLGA excluded many reads that could support expression of lncRNA genes where there was corroborating evidence for expression from reads that were not antisense to other genes. The median of reads mapped to these lncRNAs is reduced from 174 to 142 in PBMCs, 45 to 38 in liver set 1 and 70 to 57 in liver set 2 for TLGA as compared to ULGA, and the same trends are observed in each cell type (Fig 2C-D and Supplementary Fig 2A-D and Supplementary Table 1). This analysis suggests that TLGA can be used to identify lncRNAs with the highest confidence in expression but reduces the reads associated with lncRNAs containing exons antisense to other genes. On the contrary the ULGA accounts for all the possible reads mapped to lncRNA genes at the cost of potential library preparation artifacts. To this end, we combined both approaches. We utilized TLGA to define expressed lncRNAs and ULGA to account for all the reads mapped to these lncRNAs.

Two pairs of overlapping protein-coding and lncRNAs genes illustrate these scenarios in PBMCs. *SRGAP2-AS1* overlaps *SRGAP2C* in antisense (Fig 2E). The reads supporting *SRGAP2-AS1* are only within exons antisense to *SRGAP2C* (blue arrows). *SRGAP2-AS1* is defined as not expressed in TLGA and is excluded from further analysis. *HSALNG0137471* is expressed antisense to *DDX3X* (Fig 2F). In this example, there are reads supporting expression of *HSALNG0137471* in regions of exons that are not antisense to *DDX3X* exons (black vertical arrows). This lncRNA is defined as expressed in TLGA. There are additional reads mapped in ULGA closer to the 3’ end of *HSALNG0137471* that can be included to provide additional support for expression of this lncRNA.

### Read mapping and detected lncRNAs

We analyzed 8.07 billion reads in three publicly available datasets (26 samples) consisting of one PBMC dataset and two liver datasets (Table 3). We mapped all samples to GENCODE, TLGA and ULGA. Genome indices were created using Cell Ranger version 3.1.0 due to its compatibility with different versions of Cell Ranger count (material and methods). The difference in the number of reads mapped to various genomic loci between TLGA and ULGA is minimal (Supplementary Fig 3-4 & Supplementary data 1). However, we observed a significant difference between the reads mapped to GENCODE compared to both ULGA and TLGA. Across the 26 samples, we observed an increase in the reads mapped confidently to exonic regions by 1.46% in ULGA compared to GENCODE, accounting for 118.09 million reads (Supplementary Fig 3B and Supplementary data 1). These results suggest that a fraction of reads not mapped to GENCODE can be uniquely mapped to lncRNAs added with TLGA and ULGA. Furthermore, ULGA captures more lncRNA genes (expressed in at least 10 cells in a dataset) compared to GENCODE (Supplementary Table 2). GENCODE detected 5064, 4800, and 8211 lncRNAs in PBMCs, Liver set 1, and Liver set 2 compared to 25,470, 20,813, and 40,375 in ULGA. In contrast, we observed a decrease in the reads mapped confidently to the genome by 1.19% (96.26 million), intergenic regions by 1.98% (159.93 million), intronic regions by 0.69% (56 million reads), and transcriptome by 0.15% (12.09 million reads) in ULGA compared to GENCODE. The reduced reads in these categories suggest that ULGA and TLGA now captures a small number of reads defined as intergenic or intronic by GENCODE and that a small fraction of reads uniquely mapped in GENCODE are no longer uniquely mapped with the expanded annotation and are discarded (Fig 1D).

**Table 3.**
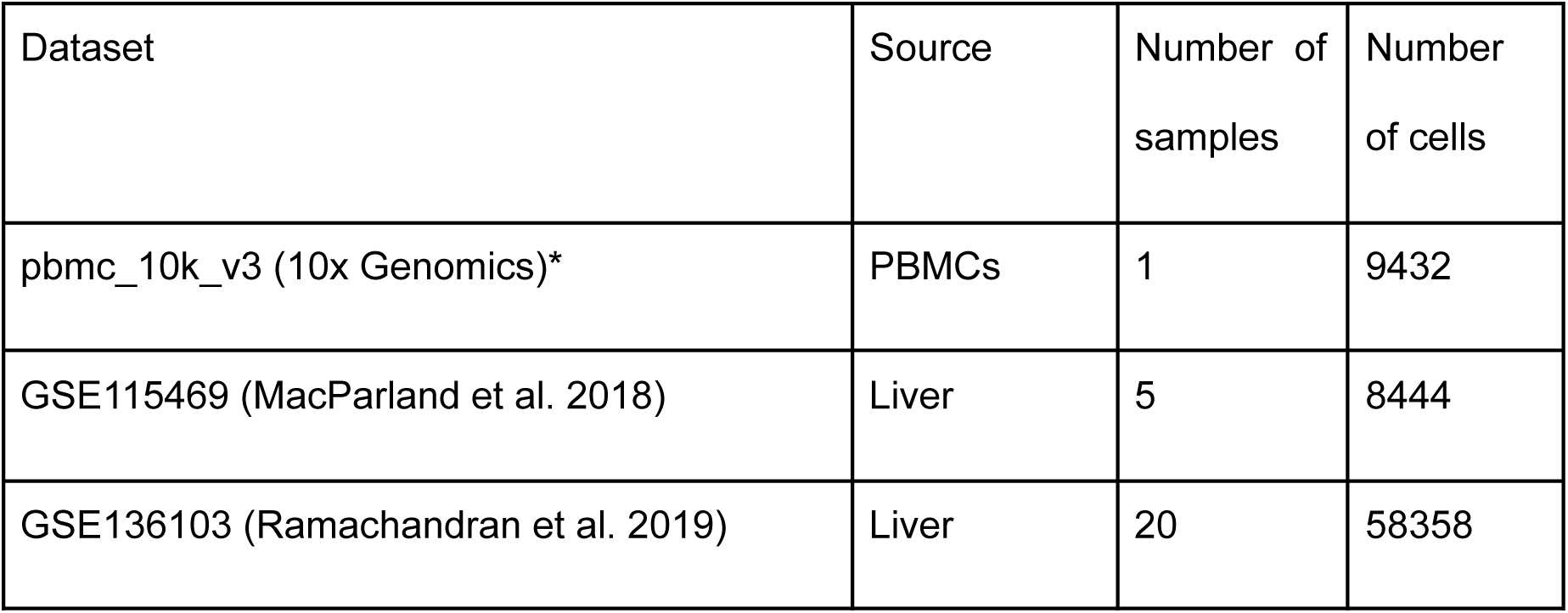
Datasets analyzed. 10x single cell RNA-seq datasets used to validate Singletrome annotation and create lncRNA cell type maps. * denotes 10k PBMCs from a Healthy Donor (v3 chemistry) Single Cell Gene Expression Dataset by Cell Ranger 3.0.0, 10x Genomics, (2018, November 19).

| Dataset | Source | Number of samples | Number of cells |
| --- | --- | --- | --- |
| pbmc_10k_v3 (10x Genomics)* | PBMCs | 1 | 9432 |
| GSE115469 (MacParland et al. 2018) | Liver | 5 | 8444 |
| GSE136103 (Ramachandran et al. 2019) | Liver | 20 | 58358 |

### Quality control of lncRNA mapping

Many lncRNAs in LncExpDB are not experimentally validated, and we next sought to define additional criteria to support lncRNA gene expression in each dataset. We assessed read distribution across the transcript body to identify lncRNA genes where 1) mapped reads exhibit 5’ bias in 3’ sequenced scRNA-seq libraries and 2) the majority of reads were mapped to a single location in the transcript, as both situations could represent library artifacts or mapping anomalies (Ma and Kingsford 2019). LncRNA genes for which all transcripts met either criteria in a dataset were excluded from further analysis in that dataset.

To obtain the read distribution across the transcript body we utilized RSeQC (Wang, Wang, and Li 2012). RSeQC scales all the transcripts to 100 bins and calculates the number of reads covering each bin position and provides the normalized coverage profile along the gene body. We modified RSeQC to obtain raw read counts (default is normalized read count to 1) for each bin (material and methods). The overall read distribution for lncRNA genes was similar to protein-coding genes in PBMCs (Fig 3A), while liver set 1 and liver set 2 showed more 5’ enrichment than protein-coding genes (Supplementary Fig 5). To assess the read distribution across the transcripts and avoid transcript length bias, we subdivided lncRNAs and protein-coding transcripts based on transcript length (Supplementary Table 3). We observed that lncRNA transcripts from 200-1000 nt in length and greater than 10,000 nt in length have very similar read distribution to protein-coding transcripts (Supplementary Fig 6-8). In contrast, the read distribution for lncRNA transcripts of length more than 1000 nt and less than 10,000 nt exhibit an increase in 5’ enrichment in 3’ sequenced scRNA-seq libraries (Fig 3B-C and Supplementary Fig 6-8).

**Figure 3.**
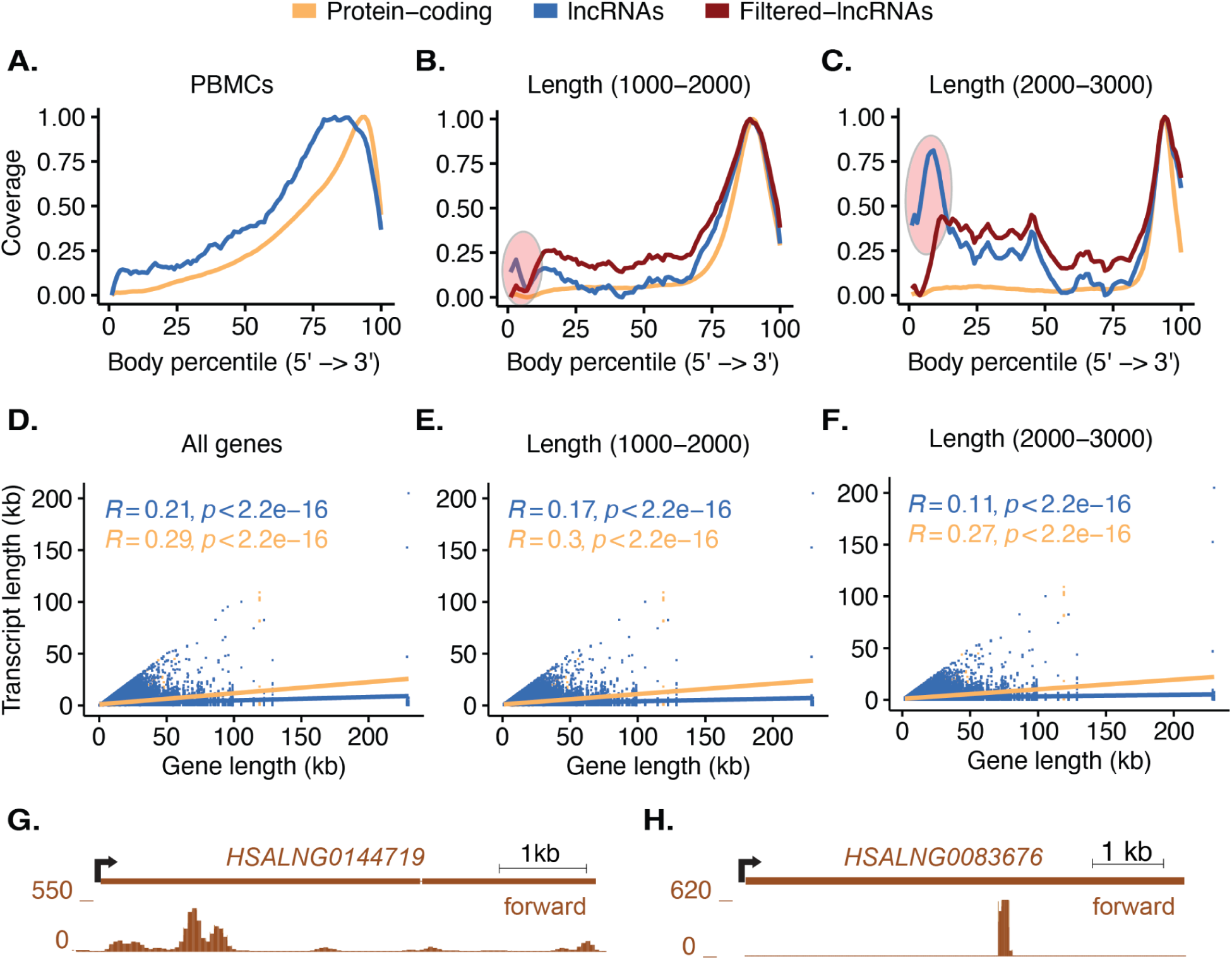
Distribution and quality control of lncRNA mapping in PBMCs. **(A)** Distribution of reads mapped across transcripts of protein-coding genes (orange) and lncRNA genes (blue). The x-axis represents RNA transcripts from 5’ to 3’ divided into 100 bins (Body percentile), and the y-axis indicates transcript coverage (0-1). The overall read distribution for lncRNA genes is similar to protein-coding genes when all transcripts are considered. **(B)** Distribution of reads mapped across transcripts from 1000-2000 nt in length. Red circle indicates an enrichment of reads in the first 10 bins of lncRNA transcripts. The transcripts responsible for this peak were identified and filtered. Filtered-lncRNAs (red line) shows the distribution of the mapped reads after removing lncRNAs that were flagged for low quality (material and methods). **(C)** Distribution of reads mapped across transcripts from 2000-3000 nt in length. Red circle indicates an enrichment of reads in the first 10 bins of lncRNA transcripts. The transcripts responsible for this peak were identified and filtered. Filtered-lncRNAs (red line) shows the distribution of the mapped reads after removing lncRNAs that were flagged for low quality (material and methods). **(D)** The correlation between transcript length (y-axis) and gene length (x-axis) is weaker for lncRNA genes (blue) than protein-coding genes (orange). Gene length (x-axis) is plotted versus transcript length (y-axis) for all lncRNAs (blue dots). The blue line indicates R, and the R value is displayed in blue. The orange line represents R for protein-coding genes, and the R value is displayed in orange. **(E)** The correlation between transcript length (y-axis) and gene length (x-axis) is plotted as in (D) for all protein-coding genes and lncRNA genes with at least one transcript with length between 1000 and 2000 nt. **(F)** The correlation between transcript length (y-axis) and gene length (x-axis) is plotted as in (D) for all protein-coding genes and lncRNA genes with at least one transcript with length between 2000 and 3000 nucleotides. **(G)** Example where reads mapped to lncRNA gene *HSALNG0144719* show that the majority of reads are from the 5’ end of the transcript and do not follow the expected distribution towards the 3’ end of the transcript. This lncRNA gene is discarded. The genomic scale is indicated on the upper right. The start site and direction of transcription are indicated by a black arrow. **(H)** Example where lncRNA gene *HSALNG0083676* has the majority of reads mapped to a single location in the transcript and this location is not at the 3’ end (last 10 bins). This lncRNA gene is discarded because this could be a library artifact or mapping anomaly.

We next assessed the variability between gene and transcript lengths and found less correlation between gene and transcript length for lncRNAs compared to protein-coding genes (Fig 3D-F & Supplementary Table 4). This finding suggested that the 5’ enrichment observed in bulk analysis of lncRNA transcripts might be explained by expression of transcripts of more variable length for a given lncRNA gene, where shorter isoforms could give the appearance of an increased fraction of 5’ reads for some lncRNA transcripts. We then evaluated 5’ bias for each lncRNA transcript (minimum transcript length 1,000 nt). In total 2445, 3065, and 4486 lncRNA genes had transcripts that were flagged for 5’ bias in PBMCs, liver set 1, and liver set 2, respectively (Fig 3G & Supplementary Table 5). Since the observed 5’ bias could be explained by more abundant shorter isoforms of an lncRNA, we discarded the lncRNA gene only if all the transcripts were flagged for 5’ bias. Using this criteria we discarded 433 lncRNA genes (5685 transcripts) in PBMCs, 488 lncRNA genes (7372 transcripts) in liver set 1, and 928 lncRNA genes (9296 transcripts) in liver set 2 (Supplementary Table 5).

Finally, we evaluated read distribution across lncRNA transcripts to identify potential library artifacts or mapping anomalies. We flagged lncRNA transcripts where reads aligned to one particular region of the full transcript (minimum transcript length 1,000 nt). If the expression of a single bin was greater than the expression of the sum of the remaining 99 bins and this single bin was not in the last 10 bins (denoting the 3’ end of the transcript), the transcript was flagged (Fig 3H). In total 606, 644, and 1084 lncRNA genes had transcripts that were flagged in the PBMCs, liver set 1, and liver set 2, respectively. We performed this analysis for all transcripts and discarded the lncRNA gene if all transcripts for a gene displayed this phenomenon. Using these criteria we discarded 67 lncRNA genes (1455 transcripts) in PBMCs, 45 lncRNA genes (1312 transcripts) in liver set 1, and 98 lncRNA genes (2271 transcripts) in liver set 2 (Supplementary Table 5).

After applying these quality control steps, we were able to retain the expression of 23,510, 19,126, and 37,507 high quality lncRNA genes in PBMCs, liver set 1, and liver set 2 (Supplementary Table 5). These lncRNAs were used for all the down-stream analyses.

### lncRNAs alone predict most clusters and cell types in single cell data

LncRNA expression can be cell-type-specific (Liu et al. 2016). We applied our new annotation to determine if we could cluster cell types based on lncRNA expression alone. We returned to scRNA-seq data for human PBMCs and liver (Table 3). We mapped scRNA-seq data using Cell Ranger (v6.0.2), and the labels for each cell were retained from the original publications. We clustered cells using data aligned to GENCODE and to Singletrome (with the previously-established filters). Despite lower expression of lncRNAs compared to protein-coding genes (Fig 2A-B & Supplementary Fig 1A-D), we created similar cell clusters based on lncRNAs alone for both PBMCs (Fig 4A-D) and liver (Supplementary Fig 9A-D & Supplementary Fig 10A-D). Clustering by lncRNAs alone showed similar results to GENCODE for most cell clusters but did shift relationships between some clusters and cell types. In PBMCs, we observed that CD4 naive and CD4 memory cells clustered more closely to CD8 naive and CD8 effector cells with lncRNAs alone compared to data aligned to GENCODE (Fig 4A and 4D). In liver set 1, we observed that hepatocytes_5 clustered closely with other hepatocytes with lncRNAs alone as compared to genome annotations containing only protein-coding genes (Supplementary Fig 9A-D). We were not able to separate sub-clusters of hepatocytes (1, 3, 6, and 15) by UMAP using lncRNAs, and these sub-clusters group closely in the original GENCODE annotation (Supplementary Fig 9A). In another example, liver sinusoidal endothelial cells (LSECs)_13 are clustered more closely with LSECs_11 and LSECs_12 with lncRNAs alone (Supplementary Fig 9D) as compared to analysis with protein-coding genes (Supplementary Fig 9A-C). For some cell types, lncRNA alone could not separate populations. For example, gamma-delta (gd)T cells_18, gd T cells_9, alpha-beta (ab) T cells, and natural killer (NK) cells could not be separated by lncRNA-only annotations (Supplementary Fig 9). It is possible that some cell types may have less diversity of lncRNA expression or produce lower levels of lncRNAs transcripts, which could reduce the ability to cluster some cell types by lncRNAs alone.

**Figure 4.**
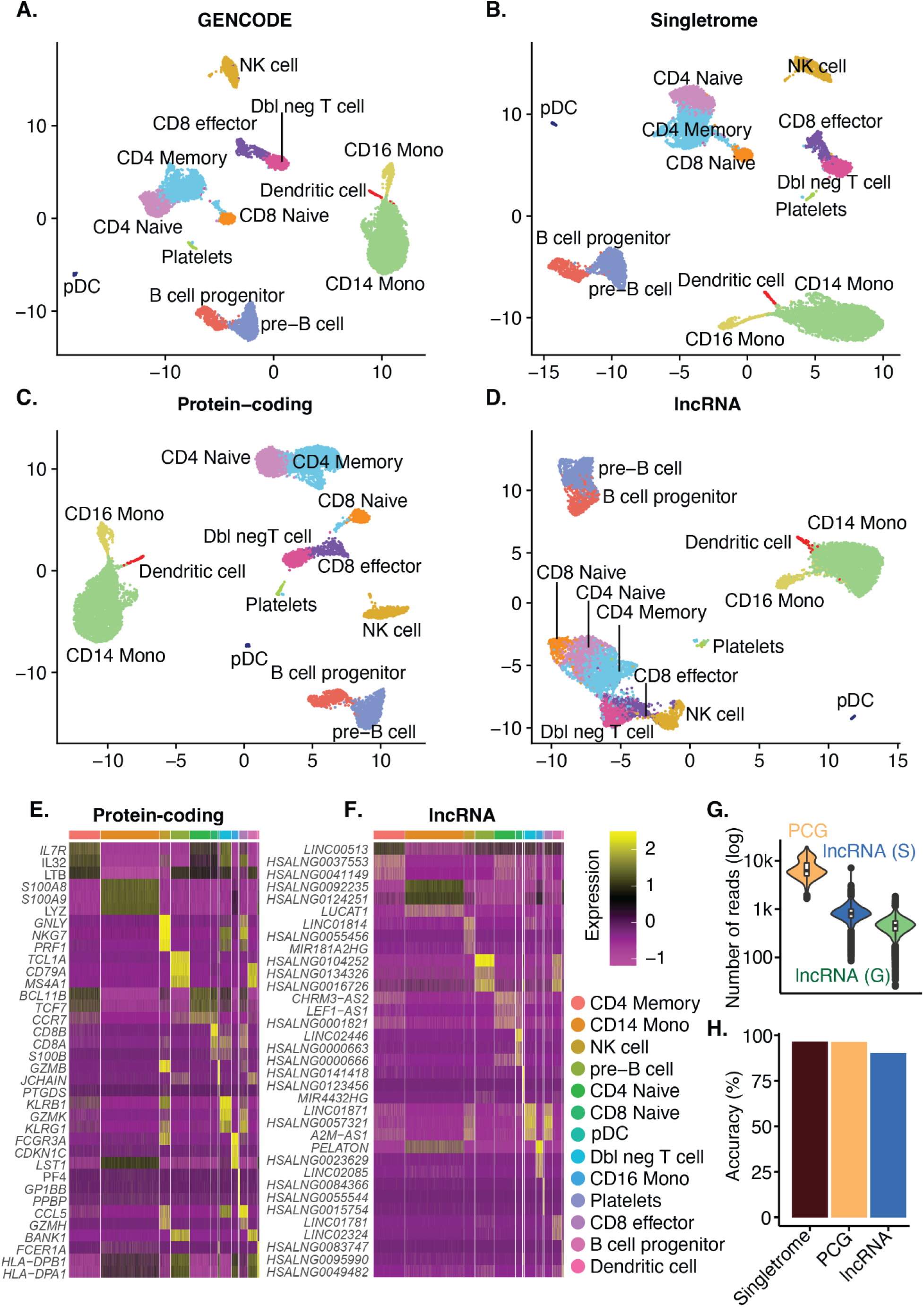
lncRNAs alone predict most clusters and cell types in single cell data. scRNA-seq data of PBMCs were mapped using annotation from **(A)** GENCODE, **(B)** Singletrome, **(C)** only protein-coding genes in Singletrome, and **(D)** only lncRNAs in Singletrome. The labels for each cell were retained from the original publications. For this analysis, Singletrome only contains lncRNAs that meet all described filters for PBMCs. **(E)** The heatmap displays the top differentially expressed protein-coding genes (y-axis) for each cell type in PBMCs. Cell types are indicated by color at the bar above the heatmap, and the key is displayed to the right. Expression level is indicated by Z-score. **(F)** The heatmap displays the top differentially expressed lncRNA genes for each cell type using the same gene expression scale as (E). Monocyte is abbreviated as Mono and double negative T cell is abbreviated as Dbl neg T cell. **(G)** The total number of mapped reads per cell (y-axis, log scale) is quantified for PCG (protein-coding genes) (orange), lncRNA (S) (lncRNA genes from Singletrome) (blue), and lncRNA (G) (lncRNA genes from GENCODE) (green) in PMBCs. **(H)** Bars showing accuracy in percentage (y-axis) for PBMC cell type prediction based on Singletrome (dark-red), PCG (protein-coding genes) from Singletrome (orange), and lncRNAs from Singletrome (blue). Receiver-operating characteristic (ROC) curves for each cell type are shown in Supplementary Fig 16.

Since lncRNAs can cluster the majority of cells by cell type, we next aimed to generate an lncRNA-based cell type marker map. We identified marker genes for each cell type relative to all other cell types based on lncRNAs and protein-coding genes in PBMCs (Fig 4E-F) and liver (Supplementary Fig 11-14). While lncRNAs are expressed at lower levels compared to protein-coding genes in all datasets (Fig 4G, Supplementary Fig 15), we were still able to identify lncRNA-based cell markers for PBMCs and liver (Supplementary data 2-4).

Clustering algorithms make assumptions about data distribution. We next trained a machine learner to determine how well lncRNAs can define cell types without the underlying statistical assumptions that are applied to clustering. In order to establish a baseline for comparing cell type predictions, we performed cell type prediction using protein-coding genes and Singletrome (containing all the protein-coding genes and quality filtered lncRNAs). We trained a gradient-boosted decision tree based classifier XGBoost (Extreme Gradient Boosting) on the expression data of protein-coding genes, lncRNAs, and the combination of both from Singletrome (material and methods). Cell type labels were retained from the original publications for PBMCs (13 cell types), liver set 1 (20 cell types), and liver set 2 (12 cell types).

We found that the overall accuracy for predicting cell types using lncRNAs was comparable to that of protein-coding genes for PBMCs (96.39% for protein-coding genes and 90.30% for lncRNAs) and liver set 2 (99.10% for protein-coding genes and 95.43% for lncRNAs) (Fig 4H, Supplementary Fig 16-17 and Supplementary data 5-6). However, liver set 1 had an accuracy of 75.48% for lncRNAs, which is considerably less than the accuracy of 93.66% for protein-coding genes (Supplementary Fig 18 Supplementary data 7). Liver set 1 splits single cell type into multiple clusters based on marker genes from GENCODE, for example it contains six cell clusters of hepatocytes, three clusters of liver sinusoidal endothelial cells (LSECs) and two cell clusters of each macrophages and gd T cells. Five out of six sub-clusters of hepatocytes are closely-associated by UMAP using the original GENCODE annotation (Supplementary Fig 9A), and two sub-clusters of each macrophages and LSECs also cluster closely using the original GENCODE annotation (Supplementary Fig 9A).

To assess the accuracy of predicting cell types rather than sub-clusters of cell types, we merged clusters within the same cell type, retaining 11 cell types in liver set 1. We were able to predict cell types with an accuracy of 98.16% using protein-coding genes, 90.40% using lncRNAs, and 98.16% using Singletrome (Supplementary Fig 19 and Supplementary data 7). These results suggest that lncRNA expression can be used to predict cell types with a similar accuracy to protein-coding genes, even though the original clustering was determined primarily by protein-coding genes.

### Long noncoding RNAs in liver fibrosis

To understand the role of lncRNAs in disease, we next analyzed scRNA-seq data of healthy and cirrhotic human liver (liver set 2, GSE136103, (Ramachandran et al. 2019). We again used cell labels from the original study and clustered the cells based on the condition (healthy and cirrhotic) using Singletrome (Supplementary Fig 20), only protein-coding genes from Singletrome (Supplementary Fig 21), and only lncRNAs from Singletrome (Fig 5A). Being able to produce similar clusters of healthy and diseased liver cell types enabled us to perform differential expression analysis of lncRNAs in healthy and cirrhotic liver by cell type.

**Figure 5.**
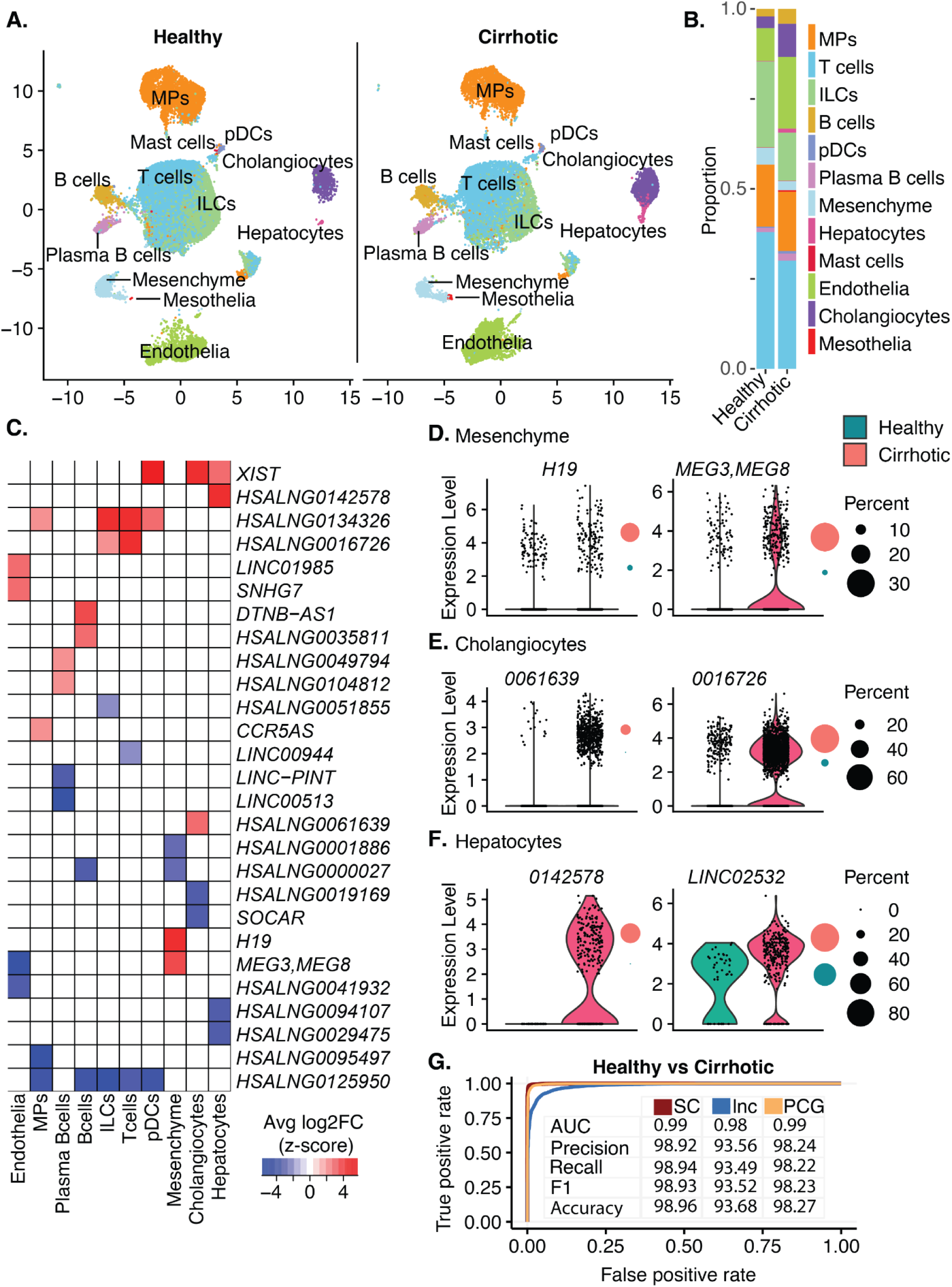
lncRNA expression predicts disease pathology. **(A)** scRNA-seq data of liver set 2 (GSE136103(Ramachandran et al. 2019)) were mapped using annotations from Singletrome. The labels for each cell were retained from the original publication. Cells were clustered based on lncRNAs from Singletrome and annotated by condition healthy (left) and cirrhotic liver (right). For this analysis, only lncRNAs that meet all the described filters in the section ‘Quality control of lncRNA mapping’ are considered. **(B)** Proportion of cells in each cell type, healthy (left) and cirrhotic liver (right). **(C)** The heatmap displays the top differentially expressed (two up- and two down-regulated) lncRNA genes (y-axis) between healthy and cirrhotic liver for each cell type. Average log2-fold is indicated by Z-score for each cell type. **(D)** Differentially regulated lncRNA expression in mesenchymal cells, **(E)** cholangiocytes and **(F)** hepatocytes. Y-axis shows the expression of the differentially regulated lncRNA gene in cells of healthy and cirrhotic liver. Circles on the right show the percentage of cells expressing the lncRNA gene in healthy (green) and cirrhotic liver (red). **(G)** Receiver-operating characteristic (ROC) curve showing true and false positive rates for healthy and cirrhotic disease prediction based on the expression of SC (all genes in Singletrome) (red), lnc (lncRNA genes) (blue) and PCG (protein-coding genes) (orange). The table shows the AUC, precision (%), recall (%), F1 (%), and accuracy (%) for healthy and cirrhotic disease prediction based on the expression of SC (all genes in Singletrome) (red), lnc (lncRNA genes) (blue) and PCG (protein-coding genes) (orange).

We detected 937 differentially expressed lncRNA genes (495 up-regulated and 442 down-regulated) between healthy and cirrhotic liver (Supplementary data 8) in cell types including mesenchymal cells, hepatocytes, cholangiocytes, endothelial cells, B cells, plasma B cells, dendritic cells (DCs), mononuclear phagocyte (MPs), innate lymphoid cells (ILCs) and T cells (padj < 0.1 and log2FC > 0.25, Supplementary data 8). We were not able to detect statistically significant differentially regulated lncRNAs in mast cells, and there were not enough mesothelial cells to perform differential expression analysis (Fig 5B).

LncRNAs induced with cirrhosis include *H19 and MEG3* in mesenchymal cells (Fig 5C-F, Supplementary data 8). *H19* has been shown to promote liver fibrosis (Wu et al. 2022; Xiao et al. 2019) *while MEG3* has been linked to both pro- and anti-fibrotic phenotypes in the liver (Zhang et al. 2017; Yu et al. 2018). While *XIST* was also enriched in specific cell types in cirrhotic samples (Fig 5C), this gene is located in the X-chromosome, and expression levels will be affected by the distribution of sexes within this dataset (cirrhotic: 2 female and 3 male; healthy: 1 female and 4 male)(material and methods). Additional lncRNAs that are not yet well-characterized were also identified as differentially expressed between cell types in healthy and cirrhotic liver (Fig 5C-F). These results show that there is sufficient read depth to identify differentially expressed lncRNAs from single cell liver datasets that could have a role in disease.

lncRNAs alone can cluster and predict cell types (Fig 4D,4H and Supplementary Fig 9D, 10D), and we were able to observe the differential expression of lncRNAs linked to cirrhosis in particular cell types. Liver pathology may be influenced by the expression of lncRNAs in affected cell types. To assess whether liver pathology follows observable rules, we trained a machine learner on lncRNA expression data of liver set 2 (GSE136103, (Ramachandran et al. 2019)) and predicted the condition (healthy or cirrhotic) of the affected target cell. Condition (healthy or cirrhotic) annotation for each cell was retained from liver set 2. We applied the XGBoost algorithm to classify healthy and cirrhotic cell types of liver using lncRNAs (material and methods). Based on lncRNA expression alone, the condition of cell types can be predicted with an accuracy of 93.68%, a precision of 93.56%, and a recall of 93.49% (Fig 5G, Supplementary data 6). In order to verify lncRNA based predictions, we trained a separate XGBoost model on the expression data of protein-coding genes, and we were able to predict the condition of the cells with an accuracy of 98.27%, a precision of 98.24%, and a recall of 98.22% (Supplementary data 6). Additionally, Singletrome was able to classify healthy and cirrhotic cells with an accuracy of 98.96%. These results suggest that it is possible for both cell type and disease pathogenicity in single cell data to be reliably predicted through analysis of lncRNA expression alone.

## Discussion

Analysis of lncRNAs has been performed in scRNA-seq data for particular sets of transcripts and cell types (Liu et al. 2016; Luo et al. 2021), but more universal lncRNA pipelines are not available. Previous approaches applied transcriptomic analysis to bulk tissue to expand known lncRNAs before evaluating expression in single cell data for the same tissue (Liu et al. 2016) or by pooling single cell data from one cell type to assemble lncRNA transcripts before analyzing expression of these transcripts at the single cell level (Luo et al. 2021). These studies demonstrate the feasibility of lncRNA analysis with single cell data and also reveal that single cell analysis can increase the sensitivity of lncRNA detection in settings where only a small population of cells express an individual lncRNA (Liu et al. 2016). Here, we sought to develop a unified analysis framework that could quantify lncRNA expression in any human scRNA-seq data.

Here, we present Singletrome, a Singularity image that integrates two GTF annotations, one containing protein-coding genes and another containing lncRNAs to generate an enhanced genome annotation. This tool provides a streamlined workflow for downstream analyses by creating BED files for further processing, running CellRanger for scRNA-seq data mapping, and merging multiple samples into a single BAM file for RSeQC quality control analysis. Additionally, it produces BigWig files for visualization. By accepting and merging two GTF files, Singletrome can accommodate a variety of annotation sources, making it a versatile tool for researchers working with different genome annotations. Our application of Singletrome focused on the LncExpDB database, but the tool is equally compatible with other lncRNA annotation sources, such as NONCODE.

While we have demonstrated Singletrome’s functionality using human genome annotations, the tool is not restricted to this species. It can be extended to any organism where both lncRNAs and protein-coding genes are defined in GTF format, providing broad applicability for genomic studies across various species. This versatility enhances the potential for Singletrome to be utilized in diverse biological contexts, facilitating high-quality analyses in genome annotation and scRNA-seq research.

We utilized Singletrome to interrogate lncRNAs in scRNA-seq data using a custom genome annotation of 110,599 genes consisting of 19,384 protein-coding genes from GENCODE and 91,215 lncRNA genes from LncExpDB (Table 1). We increased the current human lncRNA annotation by greater than five fold and benchmarked the utility of Singletrome by analyzing three publicly available 10x scRNA-seq datasets (Table 3). Mapping metrics such as reads confidently mapped to (i) exonic regions, (ii) genome, (iii) transcriptome, (iv) intronic regions, and (v) total genes detected depend on the expression of genes in each sample and can increase (with the expression of additional lncRNAs) or decrease (due to multi-mapping of reads that were originally confidently-mapped in GENCODE) with the additional lncRNA genes (Supplementary Fig 3-4 & Supplementary data 1). In most of the samples, we observed an increase in the total number of genes detected and reads confidently-mapped to exonic regions, while a decrease in the reads mapped confidently to the genome, transcriptome, intronic regions, and intergenic regions was observed in TLGA and ULGA compared to GENCODE (Supplementary Fig 4 & Supplementary data 1). The balance of reads gained by new lncRNA exons and lost by multi-mapping as a result of these exons can also fluctuate across individual samples. In the example of reads confidently mapped to exon regions (Supplementary Fig 3B), all samples show an increase for TLGA and UGLA except liver-13 and liver 25, where the loss of reads due to multi-mapping outweighs the gain of new exons.

We utilized trimmed lncRNA <u>g</u>enome <u>a</u>nnotation (TLGA) to avoid counting spurious antisense reads when defining lncRNAs that are expressed in a dataset. We then utilized an <u>u</u>ntrimmed lncRNA <u>g</u>enome <u>a</u>nnotation (ULGA) to account for all the reads mapped to lncRNAs that are defined as expressed by the TLGA (Supplementary Table 1). Using the established quality control filters, we were able to identify lncRNA genes where 1) mapped reads exhibit 5’ bias in 3’ sequenced scRNA-seq libraries and 2) the majority of reads were mapped to a single location in the transcript, as both situations could represent library artifacts or mapping anomalies(Ma and Kingsford 2019) (Fig 3G-H and Supplementary Table 5).

LncRNA expression can be cell-type-specific (Liu et al. 2016), and we found that most cell types can be clustered by lncRNAs alone (Fig 4A-D and Supplementary Fig 9-10). Clustering by lncRNAs alone was associated with less separation of clusters compared to GENCODE or lncRNAs plus protein-coding genes (Singletrome) as evidenced by the closer proximity of some clusters from the same cell type. These observations may be influenced by lower levels of lncRNA expression or more similarities in expression at the lncRNA level across similar cell types. The additional reads mapped with Singletrome did not result in the clear identification of new clusters within those clusters defined by mapping to GENCODE. However, for these analyses, cell clusters were assigned based on the originally published GENCODE annotations, and future analyses using Singletrome to perform the original clustering may provide additional resolution compared to GENCODE.

To determine cell types based on lncRNAs without the statistical assumptions, we applied the XGBoost classifier for predicting cell type using only lncRNA expression. In order to establish a baseline for comparing cell type predictions using lncRNAs, we additionally trained the XGBoost classifier on the expression data of protein-coding genes and Singletrome. We trained and predicted cell types for all the three datasets (Table 3). The classification of cells into cell types for each dataset (Table 3) was challenging due to (1) multiple cell types in each dataset and (2) imbalanced datasets (Supplementary data 5-7). There were 13 cell types in PBMCs, 20 in liver set 1, and 12 in liver set 2. Furthermore cell types were not represented equally within the dataset. For example, PBMCs have 52 platelets and 2992 CD14+ monocytes, and liver set 1 contains 37 hepatic stellate cells and over a thousand hepatocyte_1 cells. Similarly liver set 2 has 70 mast cells and over 20,000 T cells. A number of machine learning classifiers can be applied for the cell type prediction problem, such as neural networks, support vector machines, random forest and logistic regression. We selected XGBoost as it is a preferred machine learning technique for classification with imbalanced datasets against the aforementioned set of classifiers. (Hernesniemi et al. 2019; Nishio et al. 2018; Ogunleye and Wang 2020).

The overall accuracy for predicting cell types using lncRNAs was comparable to that of protein-coding genes for PBMCs (96.39% for protein-coding genes and 90.30% for lncRNAs) and liver set 2 (99.10% for protein-coding genes and 95.43% for lncRNAs). However, liver set 1 had an accuracy of 75.48% for lncRNAs, which is considerably less than the accuracy of 93.66% for protein-coding genes. Most of the mis-classifications in liver set 1 were amongst sub-clusters of the same cell type (Supplementary data 7). For example 100 cells of *Hepatocytes_3* were classified correctly, but 72 cells of *Hepatocytes_3* are classified as *Hepatocytes_1.* These results indicate that the cells in each subcluster still contain many of the same lncRNAs. When we collapsed cell sub-clusters into a single cell type, we predicted liver set 1 cell types with an accuracy of 98.16% for protein-coding and 90.40% for lncRNAs (Supplementary data 7). These results suggest that lncRNA expression can be used to predict cell types with a similar accuracy to protein-coding genes. However, the prediction accuracy drops when separating sub-clusters of the same cell type. In these cases it is not yet clear whether lncRNAs are slightly less able to predict cell type, or the difference in prediction between lncRNAs and protein-coding genes reflects an inherent bias towards the protein-coding genes in the original cell type labeling.

The ability to cluster and predict most cell types using lncRNA expression alone, demonstrates the depth and diversity of lncRNA transcripts detected in single cell data. The overall goal of Singletrome is to increase the depth of annotations of single cell data and to define differentially expressed lncRNA genes that may regulate disease processes. Comparing cells from healthy and cirrhotic liver (liver set 2), we were able to identify 937 differentially expressed lncRNAs. *H19* and *MEG3* are both linked to liver fibrosis (Xiao et al. 2019; Zhang et al. 2017; Yu et al. 2018) and were identified in our analysis. This suggests that other lncRNAs with similar patterns of expression (Fig 5C-F and Supplementary data 8) may also have activity in liver fibrosis. Our analysis was based on currently available data for healthy and cirrhotic liver. The data includes five cirrhotic livers with different causes of cirrhosis. As liver datasets expand in the future to include additional replicates with cirrhosis from multiple sources of injury and different stages along disease progression, the statistical power will increase to allow identification of additional differentially expressed lncRNAs across all conditions.

lncRNAs alone can cluster and predict cell types (Fig4D, 4H and Supplementary Fig 9D, 10D, 16-19), and we were able to identify differential regulation of lncRNAs linked to cirrhosis in particular cell types. Machine learning also demonstrated the ability of lncRNAs to predict disease (Fig 5G). These analyses demonstrates that lncRNA expression changes significantly in disease and provides further support to suggest that lncRNAs, in addition to protein-coding genes, can serve as biomarkers and mechanistic drivers of disease (Nath et al. 2019; Delás and Hannon 2017; Bolha, Ravnik-Glavač, and Glavač 2017).

The Human Cell Atlas has now mapped more than a million individual cells across 33 organs of the human body (Suo et al. 2022; Domínguez Conde et al. 2022). The focus of these analyses has understandably been on protein-coding genes. This comprehensive genome annotation optimized for scRNA-seq data can now be applied to existing and future single cell data sets to promote the development of an atlas of human lncRNAs in health and disease.

## Material and methods

### Container environment and dependencies

Singletrome is provided as a Singularity image, which requires an apptainer. Users may download the prebuilt SIF container or rebuild it using the provided build.apptainer script (section code availability). The container includes essential bioinformatics tools installed via Miniconda (e.g., BEDTools, RSeQC, DeepTools, and CellRanger). The Singletrome pipeline, executed via the Singletrome.py script, integrates two GTF annotations, one containing protein-coding genes and another containing lncRNAs. It automates tasks such as downloading and preprocessing GTF files, analyzing exon overlaps using BEDTools, merging annotations into a final GTF file, and builds a genome index for CellRanger. Furthermore, if multiple samples are analyzed and mapped using the run_cellranger.py script, it will generate a merged BAM file for RSeQC quality control analysis and a BigWig file for visualization.

#### Genome indices

We downloaded the human reference genome index from 10x Genomics https://cf.10xgenomics.com/supp/cell-exp/refdata-gex-GRCh38-2020-A.tar.gz, which includes genes from different biotypes (lncRNA, protein_coding, IG_V_pseudogene, IG_V_gene, IG_C_gene, IG_J_gene, TR_C_gene, TR_J_gene, TR_V_gene, TR_V_pseudogene, TR_D_gene, IG_C_pseudogene, TR_J_pseudogene, IG_J_pseudogene, IG_D_gene) as shown in Supplementary Table 6 along with the number of genes for each biotype. We termed this genome annotation as GENCODE (used by Cell Ranger) in the manuscript. For evaluating protein-coding and lncRNAs exonic overlap in the GENCODE annotation, we used the same strategy and script from 10x Genomics with protein_coding and lncRNA as the biotype patterns respectively. In brief, we obtained 19,384 protein-coding genes with GENCODE v32 filtering for ‘protein_coding’ as the ‘gene_type’ and ‘transcript_type’. We additionally filtered transcripts with tags such as ‘readthrough_transcript’ and ‘PAR’. We obtained 16,849 long noncoding RNAs filtering GENCODE v32 for ‘lncRNA’ as the ‘gene_type’ and ‘transcript_type’. We additionally filtered transcripts with tags such as ‘readthrough_transcript’ and ‘PAR’. For TLGA and ULGA genome indices, we downloaded the human lncRNA genome annotation file from ftp://download.big.ac.cn/lncexpdb/0-ReferenceGeneModel/1-GTFFiles/LncExpDB_OnlyLnc.tar.gz

We removed 8 genes (HSALNG0056858, HSALNG0059740, HSALNG0078365, HSALNG0092690, HSALNG009306, HSALNG0089130, HSALNG0089954 and HSALNG0095105) where we found invalid exons in the transcript or exons of transcripts were not stored in ascending order. To create the TLGA and ULGA genome indices, we included the protein-coding genes obtained from the GENCODE with the inhouse created genome annotation file (see section ‘Expanding lncRNA annotations in single cell analysis’), and created the genome indices using the bash script available at 10x Genomics website (https://support.10xgenomics.com/single-cell-gene-expression/software/release-notes/buildhg19_3.0.0). For all the genome indices, the human reference sequence for GRCh38 was downloaded from http://ftp.ensembl.org/pub/release-98/fasta/homo_sapiens/dna/Homo_sapiens.GRCh38.dna.pri <u>mary_assembly.fa.gz</u>. Genome indices were created using Cell Ranger version 3.1.0 due its compatibility with all the versions (3.1 to 6.0 as of the current work) of count pipelines and older v1 chemistry versions of Cell Ranger count (Supplementary Table 7). Using Cell Ranger version 3.1.0 mkref will help to analyze scRNA-seq data generated with the older v1 chemistry version.

#### Data

We analyzed three publicly available 10x scRNA-seq datasets consisting of 26 samples (Table 3) with the most widely used genome annotation for scRNA-seq analysis (GENCODE) and our custom genome annotations (TLGA and ULGA).

#### Gene expression

Cell Ranger count version 6.0.2 was used with the default parameters for all the genome versions to obtain gene expression count matrix.

#### lncRNA quality filter

To compute the gene body coverage for each dataset (PBMCs, liver set 1 and liver set 2) we utilized RSeQC (Wang, Wang, and Li 2012). The program was used to check if reads coverage was uniform and if there was any 5’ or 3’ end bias or if majority of the reads are mapped to one location (single bin) in the transcript. RSeQC scales all the transcripts to 100 bins and calculates the number of reads covering each bin position and provides the normalized coverage profile along the gene body. We modified the RSeQC geneBody_coverage.py script to obtain raw read counts (default is normalized read count to 1) for each bin. To assess the read distribution across the gene body and avoid transcript length bias, we subdivided lncRNAs and protein-coding transcripts based on transcript length. Gene and transcript length were calculated using R package GenomicFeatures version 1.46.1 (Supplementary Table 3). The input for the program is an indexed BAM file and gene model in BED format. Gene models were created for protein-coding genes from GENCODE and lncRNAs from Singletrome. We assessed read distribution across the transcript body to identify lncRNA genes where 1) mapped reads exhibit 5’ bias in 3’ sequenced scRNA-seq libraries and 2) the majority of reads were mapped to a single location in the transcript, as both situations could represent library artifacts or mapping anomalies (Ma and Kingsford 2019). LncRNA genes for which all transcripts met either criteria in a dataset were excluded from further analysis in that dataset. LncRNAs that passed these filtering steps were used for all the downstream analysis such as cell type clustering, cell type prediction, differential expression in healthy and cirrhotic liver and disease prediction.

#### Cell type clustering

We used Seurat version 4.0.6 for analyzing all the gene expression matrices for all the datasets (Table 3). We retained cell type labels from the original publications. We matched the barcodes from our mapping to the original publication barcodes to obtain the cell type labels. In all the analyses, the Singletrome count matrix was subsetted for protein-coding genes and lncRNAs to cluster cell types. Since we used the cell labels from the original publications (Table 3), we discarded all the other cells that were not labeled (assigned cell type identity) in the original publication.

#### Cell type markers identification

We used Seurat version 4.0.6 for the identification of cell type markers in all the datasets (Table 3). To identify cell type markers based on lncRNA and protein-coding genes, gene expression count matrices obtained from Singletrome mapping were split into protein-coding and lncRNA genes for each dataset. We used FindAllMarkers function from Seurat to find markers (differentially expressed genes) for each of the cell types in a dataset. We retained only those genes with a log-transformed fold change of at least 0.25 and expression in at least 25% of cells in the cluster under comparison.

#### Cell type prediction using machine learning

We trained a XGBoost classifier (version 1.6.2) on the expression data of protein-coding genes, lncRNAs, and the combination of both in Singletrome to predict cell types. Cell type labels were retained from the original publications (Table 3). We opted for XGBoost, as it is a preferential model for the imbalanced data and some cell types were under-represented in the datasets (Table 3 and Supplementary data 5-7). Expression data for each model (protein-coding, lncRNAs and Singletrome) was split into a training set (80%) and test set (20%). The model was trained using 80% of the data and evaluated using the remaining 20% of the data for each dataset (Table 3). To find the optimal parameters for the model, we used RandomizedSearchCV. The resultant optimal parameters for cell type classification were n_estimators : 25, max_depth : 25 and tree_method : ’hist’. Measurements of the model performance such as accuracy, recall, precision, f1, specificity, AUC are reported for each model for all the datasets (Supplementary data 5-7).

#### Differential expression analysis

We used Seurat version 4.0.6 to perform differential expression analysis between healthy and cirrhotic liver for liver set 2. The gene expression count matrix obtained from Singletrome mapping was split into protein-coding and lncRNA genes. Differential expression analysis was performed separately for Singletrome, protein-coding genes and lncRNA genes. We used FindMarkers function from Seurat to identify the differentially expressed genes between healthy and cirrhotic liver for each cell type. We filtered differentially expressed genes (protein-coding and lncRNAs) for padj-value less than 0.1 and log2FC more than 0.25 in either direction.

#### Disease (cirrhosis) prediction using machine learning

We trained XGBoost classifier (version 1.6.2) on the expression data of protein-coding genes, lncRNAs, and the combination of both in Singletrome to predict the condition (healthy or cirrhotic) of the cell in liver set 2. Condition (healthy or cirrhotic) labels were retained from the original publication (liver set 2). RandomizedSearchCV technique was used to identify the optimum values of various parameters for the model. The optimum values obtained for various parameters were n_estimators: 400, max_depth: 25, subsample: 0.75, and tree_method:’hist’. Expression data for each model (protein-coding, lncRNAs and Singletrome) was split into a training set (80%) and test set (20%). The model was trained using 80% of the data and evaluated using the remaining 20% of the data. Measurements of the model performance such as accuracy, recall, precision, f1, specificity, AUC are reported for each model (Supplementary data 6).

#### Sex Determination of Liver Set 2 Samples

The original publication (Ramachandran et al. 2019) and GEO dataset (GSE136103) did not provide sex information for the Liver Set 2 samples. To infer the sex of each sample, we analyzed the expression of X- and Y-chromosome-associated genes, including *RPS4Y1*, *XIST*, and *ZFX*. High expression of Y-linked genes (*RPS4Y1*) was used as an indicator of male samples, whereas the presence of *XIST*, an X-chromosome gene involved in X-inactivation, suggested female samples. Based on this approach, we inferred that the cirrhotic group consisted of two female and three male samples, while the healthy group included one female and four male samples.

## Data availability

All the datasets (Table 3) used in this study are publicly available. The PBMCs dataset was obtained from the 10x Genomics platform “10k PBMCs from a Healthy Donor (v3 chemistry) Single Cell Gene Expression Dataset by Cell Ranger 3.0.0, 10x Genomics, (2018, November 19)”. The previously-published datasets from the Gene Expression Omnibus (GEO) used in this study are GSE115469 and GSE136103.

## Code availability

Python, R and Bash Scripts for data processing are available through https://github.com/RAZA-UR-RAHMAN/Singletrome.

## Acknowledgments

The authors thank Kate Jeffrey for helpful discussion. A.C.M. was supported by the Chan Zuckerberg Initiative, Pew Biomedical Scholars Program, and NIH/NIDDK R01DK116999. This publication is part of the Human Cell Atlas - www.humancellatlas.org/publications.

## Author contributions

R.R and A.C.M. conceived and designed the study. Computational analysis was performed by R.R. Z.L developed and tested the singularity image with input from R.R. I.A. and R.R. designed the cell type and disease prediction analysis and I.A implemented the Xgboost models. R.S and A.S assisted with the analysis of the differentially expressed lncRNAs in liver fibrosis. The manuscript was written by R.R. and A.C.M with input from all other authors.

## Supplementary Figures

**Supplementary Figure 1.**
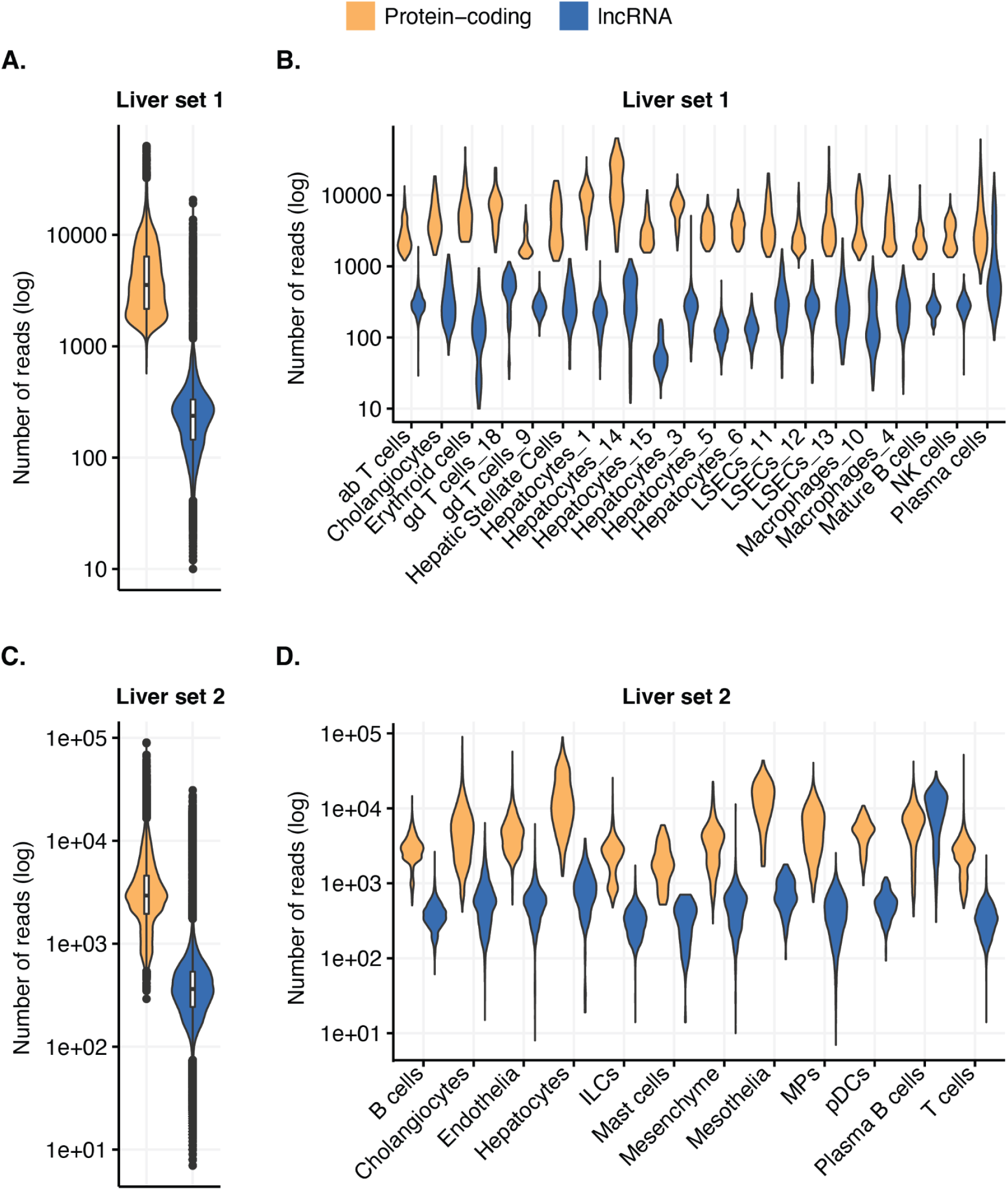
Distribution of transcripts in bulk and by cell type in liver. LncRNAs (blue) are expressed at lower levels than protein-coding genes (orange) in liver set 1 **(A),** all cell types of liver set 1 **(B),** liver set 2 **(C),** and all cell types of liver set 2 **(D)**. Reads aligned to lncRNAs and protein-coding genes are shown in log scale (y-axis). Cell types (x-axis) are determined by the original description from liver set 1 (GSE115469 (MacParland et al. 2018)) and liver set 2 (GSE136103 (Ramachandran et al. 2019)).

**Supplementary Figure 2.**
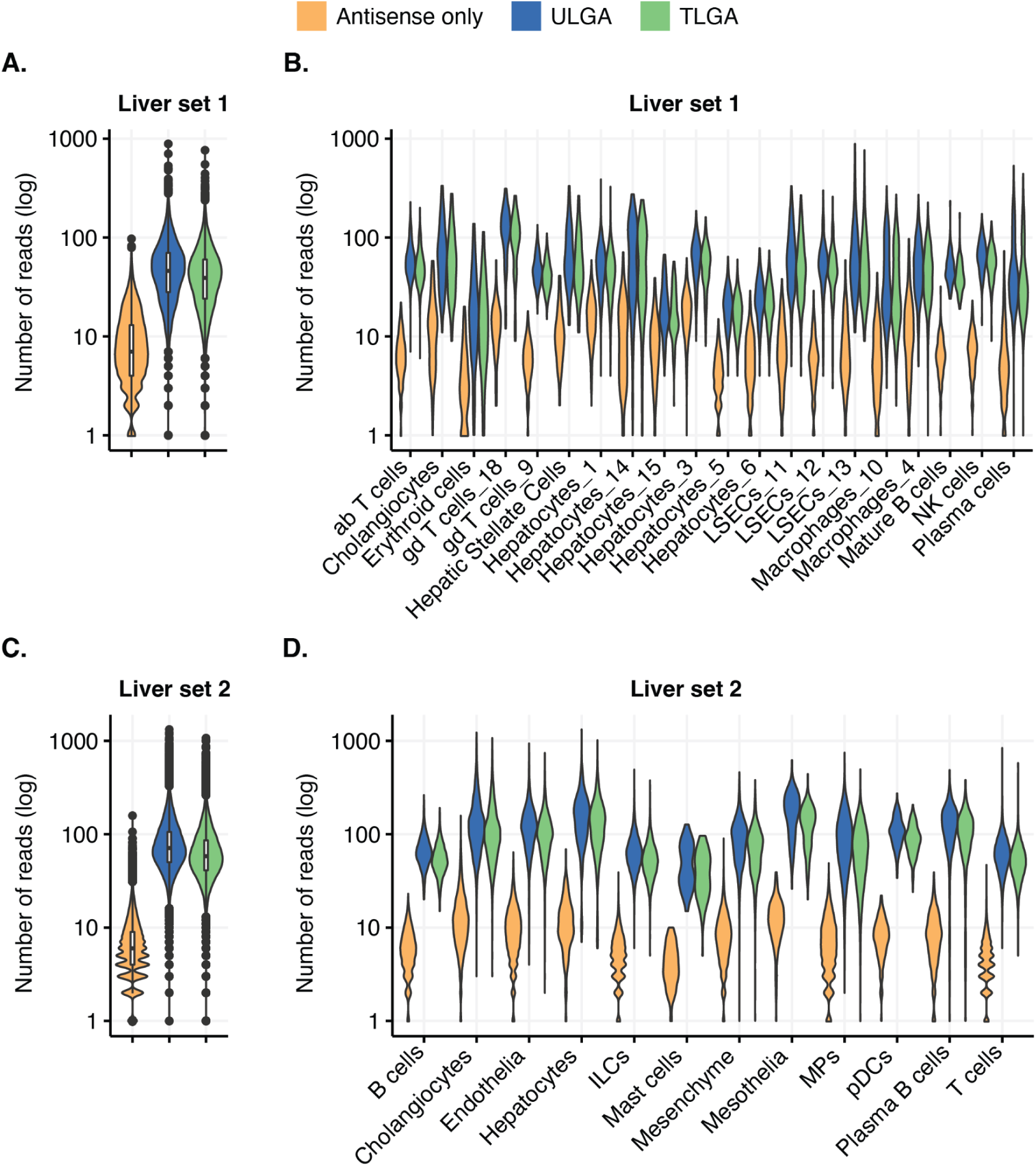
Loss of lncRNA expression due to trimming in bulk and by cell type in liver. Expression of lncRNAs that overlap protein-coding genes in the antisense direction. Expression of lncRNAs in non-overlapping regions [TLGA (green)], expression of lncRNAs in overlapping and non-overlapping regions [ULGA (blue)], and lncRNA exons that are expressed only in regions antisense to protein-coding exons [antisense only (orange)] are displayed (y-axis). The Y-axis shows the number of reads in log scale. TLGA (green) identifies lncRNAs with the highest confidence in expression but reduces the reads associated with lncRNAs as compared to ULGA (blue) in liver set 1 **(A),** all cell types of liver set 1 **(B),** liver set 2 **(C)** and all cell types of liver set 2 **(D)**. Cell types (x-axis) are determined by the original description from liver set 1 (GSE115469 (MacParland et al. 2018)) and liver set 2 (GSE136103 (Ramachandran et al. 2019)).

**Supplementary Figure 3.**
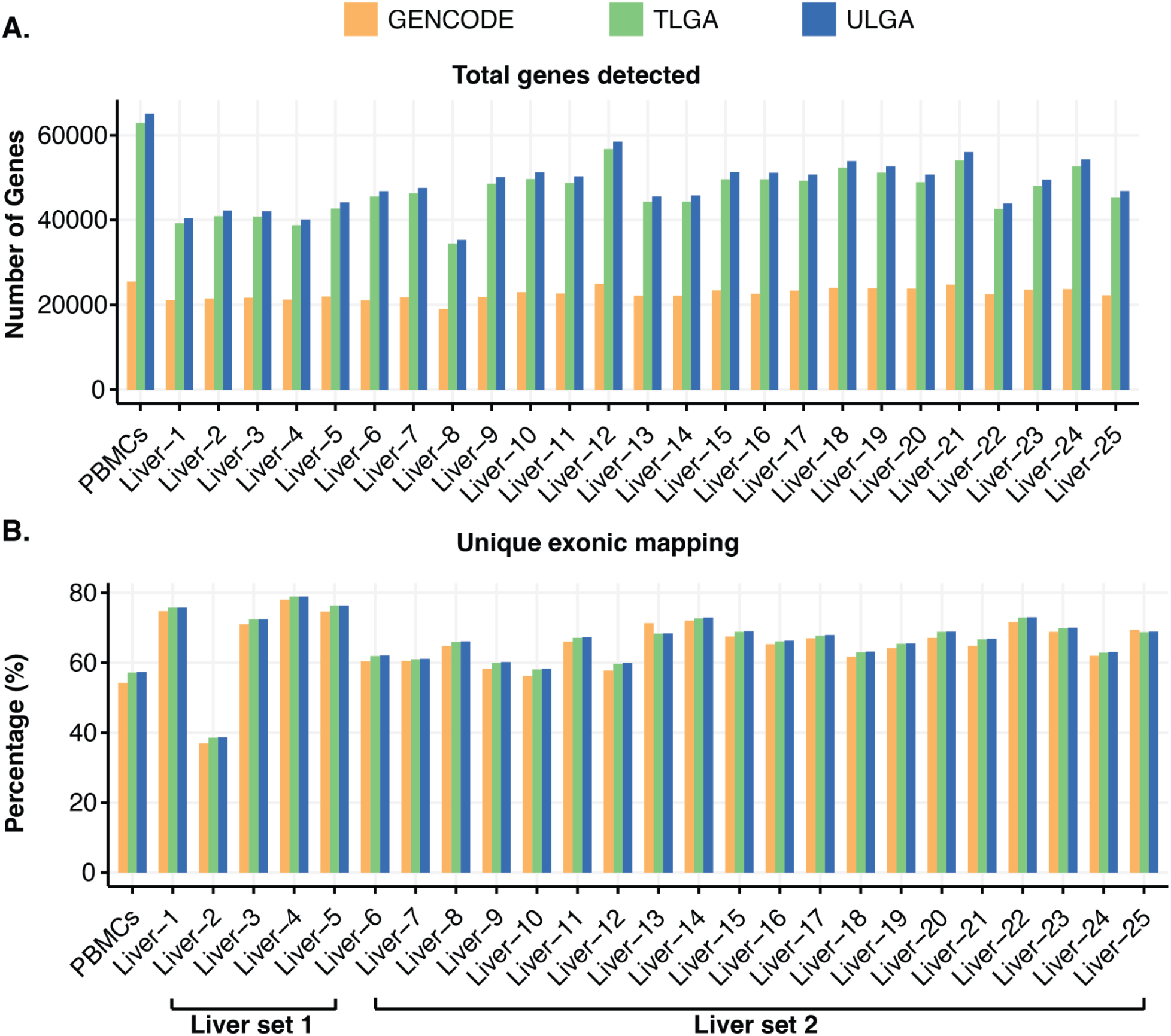
Comparison of total genes detected and unique exonic mapping with GENCODE, TLGA, and ULGA annotations. **(A)** The total number of genes mapped (y-axis) increases in TLGA and ULGA compared to GENCODE. Analysis was performed for PBMCs (10x genomics) and liver set 1 (liver 1-5) (MacParland et al. 2018) and liver set 2 (liver 6-25) (Ramachandran et al. 2019). **(B)** The percentage of reads mapped uniquely to exons in TLGA and ULGA compared to GENCODE in PBMCs, samples from liver set 1, and liver set 2.

**Supplementary Figure 4.**
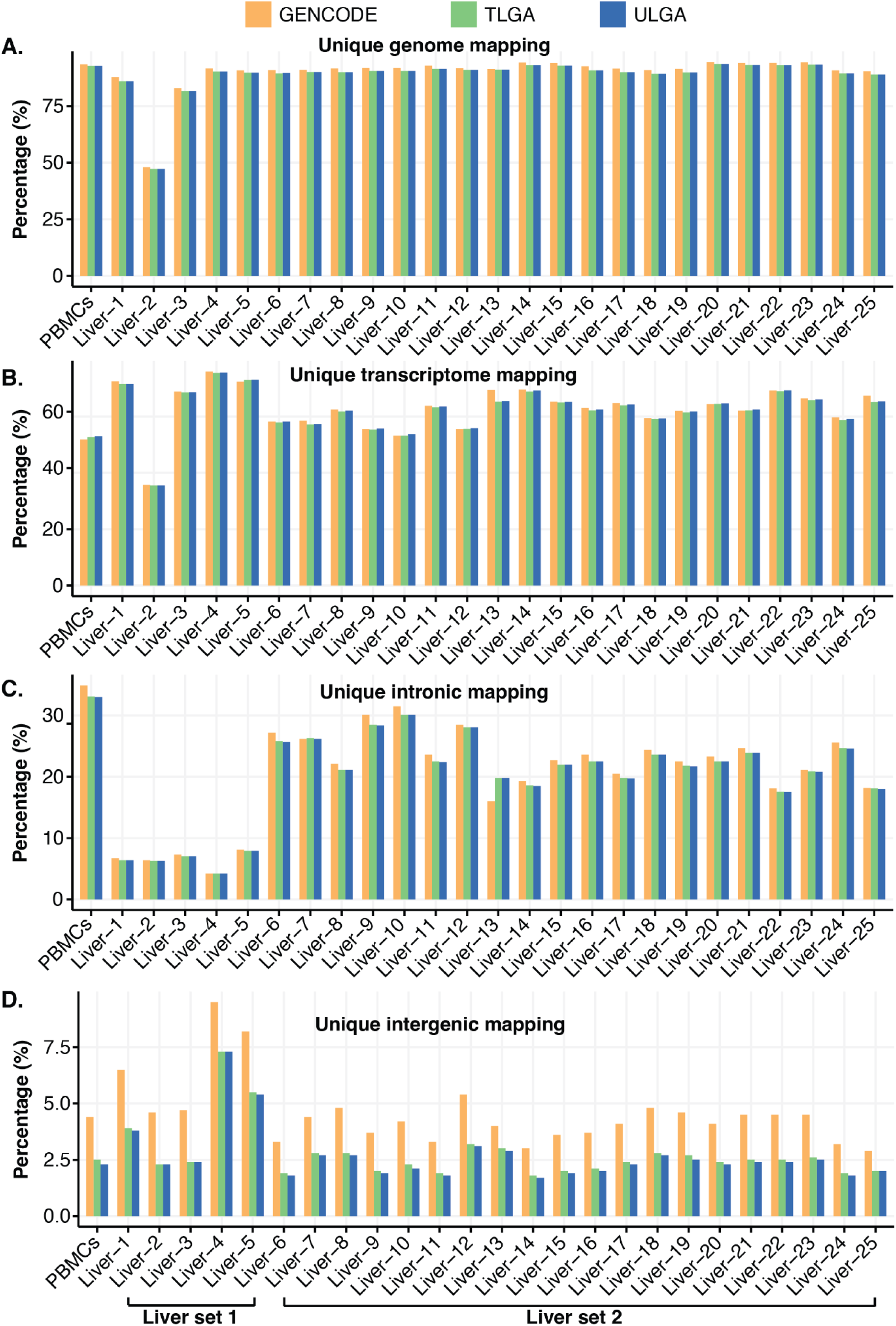
Comparison of reads uniquely mapped to genome, transcriptome, intronic regions and intergenic regions with GENCODE, TLGA, and ULGA annotations. Y-axis shows percentage of reads mapped uniquely to **(A)** genome, **(B)** transcriptome, **(C)**, intronic regions and **(D)** intergenic regions with TLGA and ULGA compared to GENCODE in PBMCs, samples from liver set 1 (liver 1-5) (MacParland et al. 2018) and liver set 2 (liver 6-25) (Ramachandran et al. 2019).

**Supplementary Figure 5.**
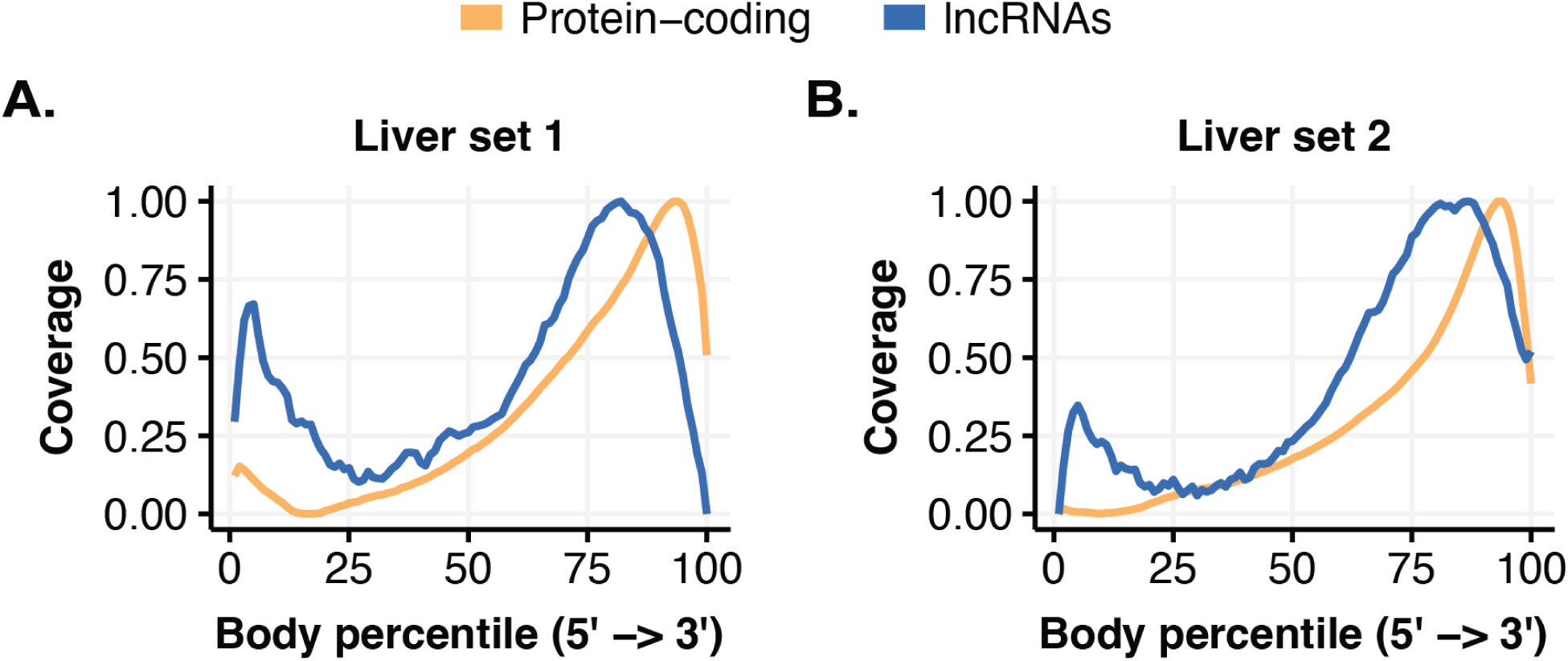
Distribution of the mapped reads across the transcripts in liver. Distribution of reads mapped across transcripts of protein-coding genes (orange) and lncRNA genes (blue) in liver. The x-axis represents RNA transcripts from 5’ to 3’ divided into 100 bins (Body percentile), and the y-axis indicates transcript coverage (0-1). **(A)** Expressed protein-coding genes (orange) and lncRNAs (blue) in liver set 1. **(B)** Expressed protein-coding genes (orange) and lncRNAs (blue) in liver set 2. In both liver set 1 and liver set 2 lncRNAs (blue) transcripts indicate an enrichment of reads at the 5’ end of lncRNA transcripts.

**Supplementary Figure 6.**
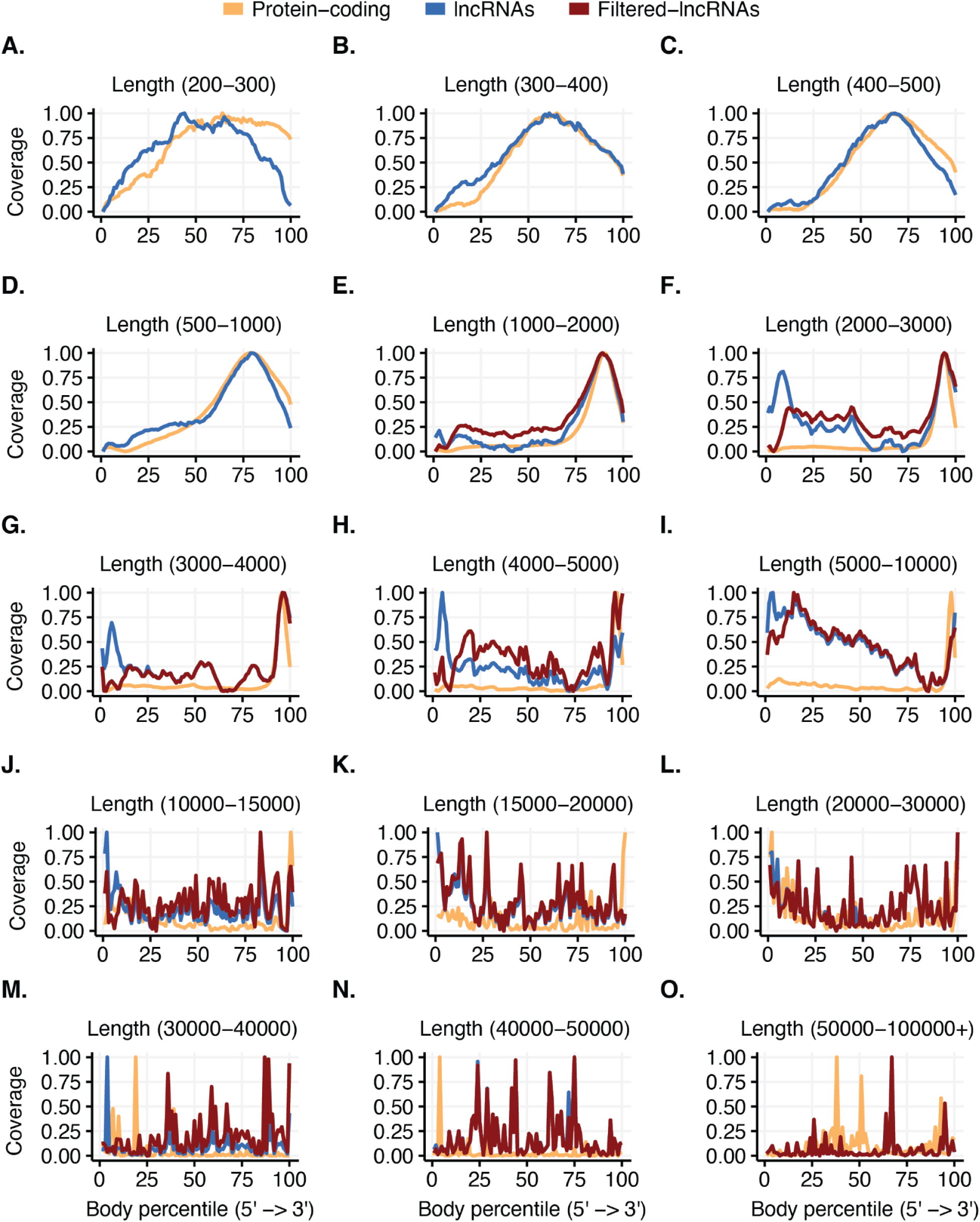
Distribution of lncRNA mapping in PBMCs across transcripts of different lengths. **(A-O)**, distribution of reads mapped across transcripts of protein-coding genes (orange) and lncRNA genes (blue) across different lengths of transcripts. Minimum and maximum length of the transcripts for each panel are shown at the top in parenthesis. The x-axis represents RNA transcripts from 5’ to 3’ divided into 100 bins (Body percentile), and the y-axis indicates transcript coverage (0-1). lncRNA transcripts of 1000 or more nucleotides **(E-O)** were filtered, if reads mapped to a transcript indicate an enrichment of reads in the first 10 bins of lncRNA transcript, or if the majority of reads were mapped to a single location (1 bin) in the transcript and that location is in the first 90 bins. Filtered-lncRNAs (red line) shows the distribution of the mapped reads after removing lncRNAs that were flagged for low quality (material and methods).

**Supplementary Figure 7.**
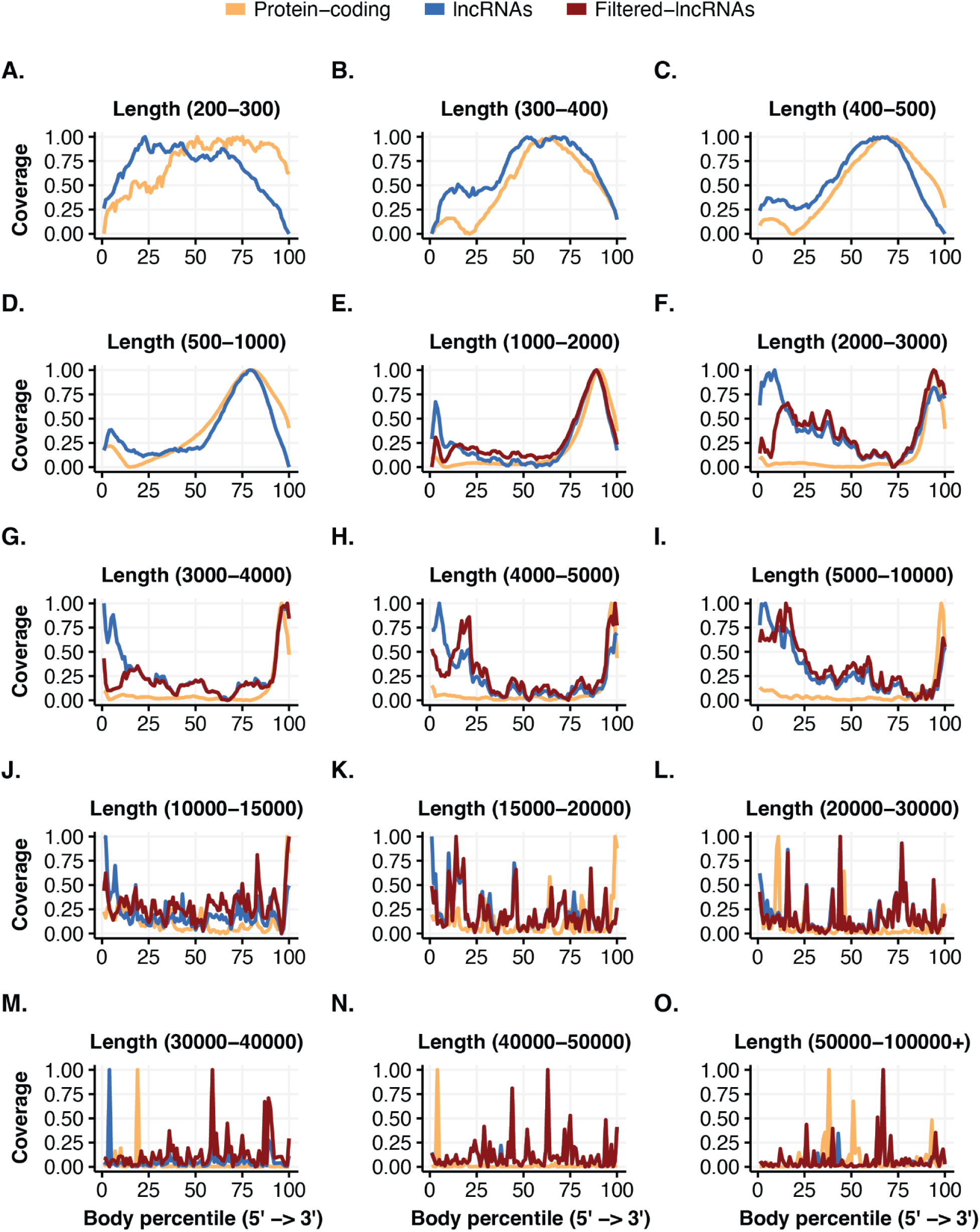
Distribution of lncRNA mapping in liver set 1 across transcripts of different lengths. **(A-O)**, distribution of reads mapped across transcripts of protein-coding genes (orange) and lncRNA genes (blue) across different lengths of transcripts. Minimum and maximum length of the transcripts for each panel are shown at the top in parenthesis. The x-axis represents RNA transcripts from 5’ to 3’ divided into 100 bins (Body percentile), and the y-axis indicates transcript coverage (0-1). lncRNA transcripts of 1000 or more nucleotides **(E-O)** were filtered, if reads mapped to a transcript indicate an enrichment of reads in the first 10 bins of lncRNA transcript, or if the majority of reads were mapped to a single location (1 bin) in the transcript and that location is in the first 90 bins. Filtered-lncRNAs (red line) shows the distribution of the mapped reads after removing lncRNAs that were flagged for low quality (material and methods).

**Supplementary Figure 8.**
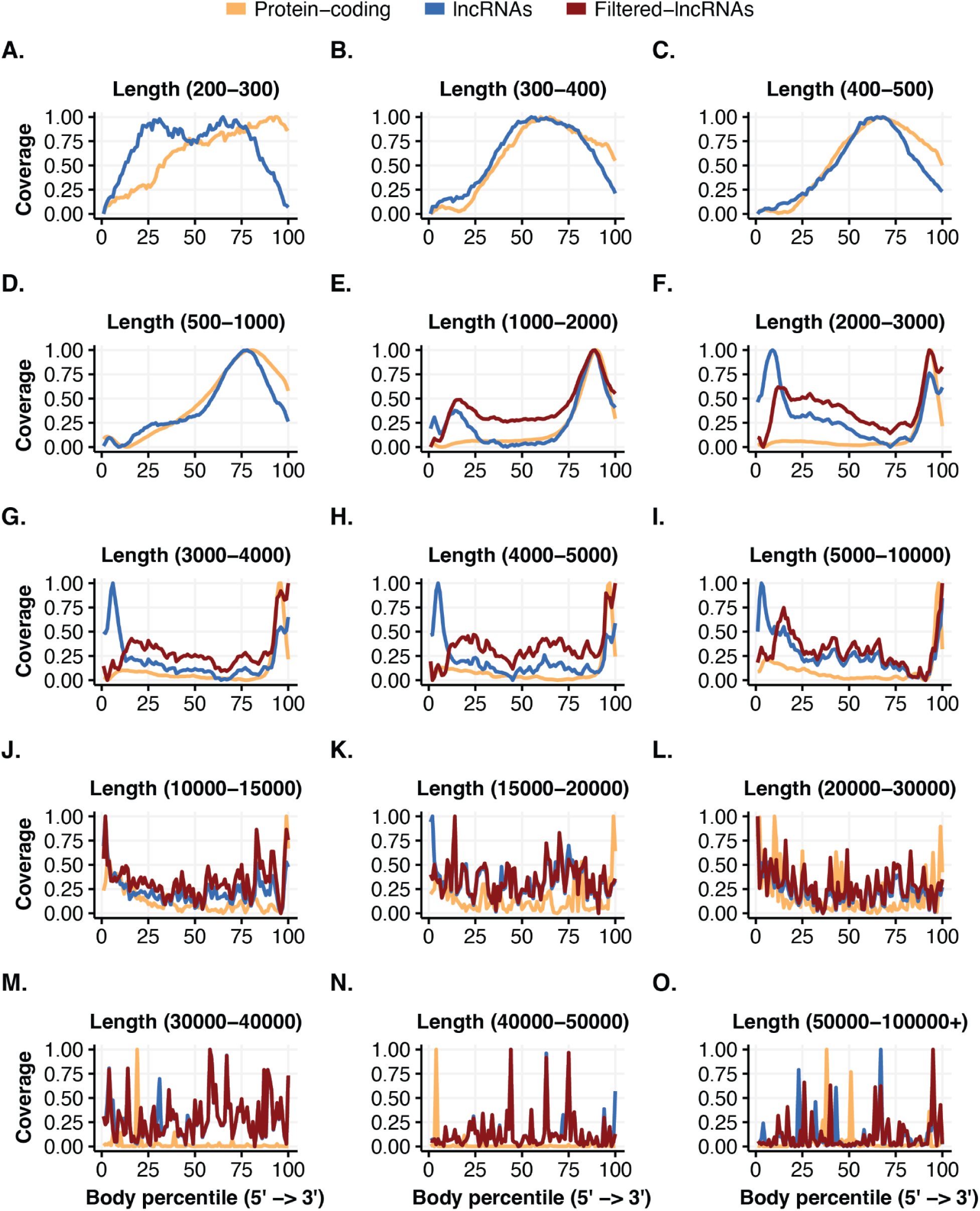
Distribution of lncRNA mapping in liver set 2 across transcripts of different lengths. **(A-O)**, distribution of reads mapped across transcripts of protein-coding genes (orange) and lncRNA genes (blue) across different lengths of transcripts. Minimum and maximum length of the transcripts for each panel are shown at the top in parenthesis. The x-axis represents RNA transcripts from 5’ to 3’ divided into 100 bins (Body percentile), and the y-axis indicates transcript coverage (0-1). lncRNA transcripts of 1000 or more nucleotides **(E-O)** were filtered, if reads mapped to a transcript indicate an enrichment of reads in the first 10 bins of lncRNA transcript, or if the majority of reads were mapped to a single location (1 bin) in the transcript and that location is in the first 90 bins. Filtered-lncRNAs (red line) shows the distribution of the mapped reads after removing lncRNAs that were flagged for low quality (material and methods).

**Supplementary Figure 9.**
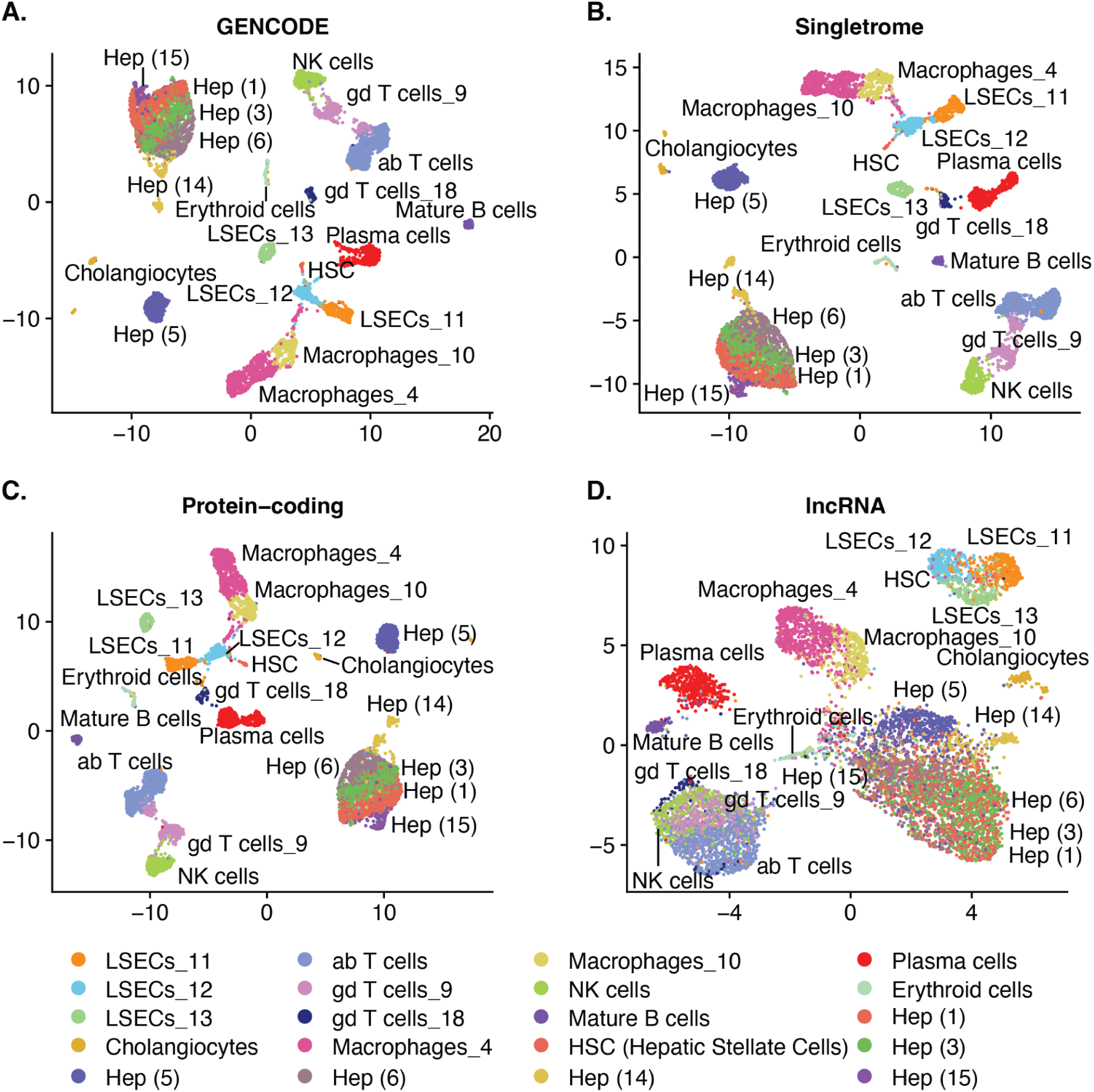
lncRNAs alone predict most clusters and cell types in single cell data of liver set 1. scRNA-seq data of liver set 1 were mapped using annotation from **(A)** GENCODE, **(B)** Singletrome, **(C)** only protein-coding genes in Singletrome, and **(D)** only lncRNAs in Singletrome. The labels for each cell were retained from the original publication. For this analysis, Singletrome only contains lncRNAs that meet all filters developed with analysis of PBMCs and applied to data from liver set 1. Hepatocytes are abbreviated as Hep and hepatic stellate cells are abbreviated as HSC.

**Supplementary Figure 10.**
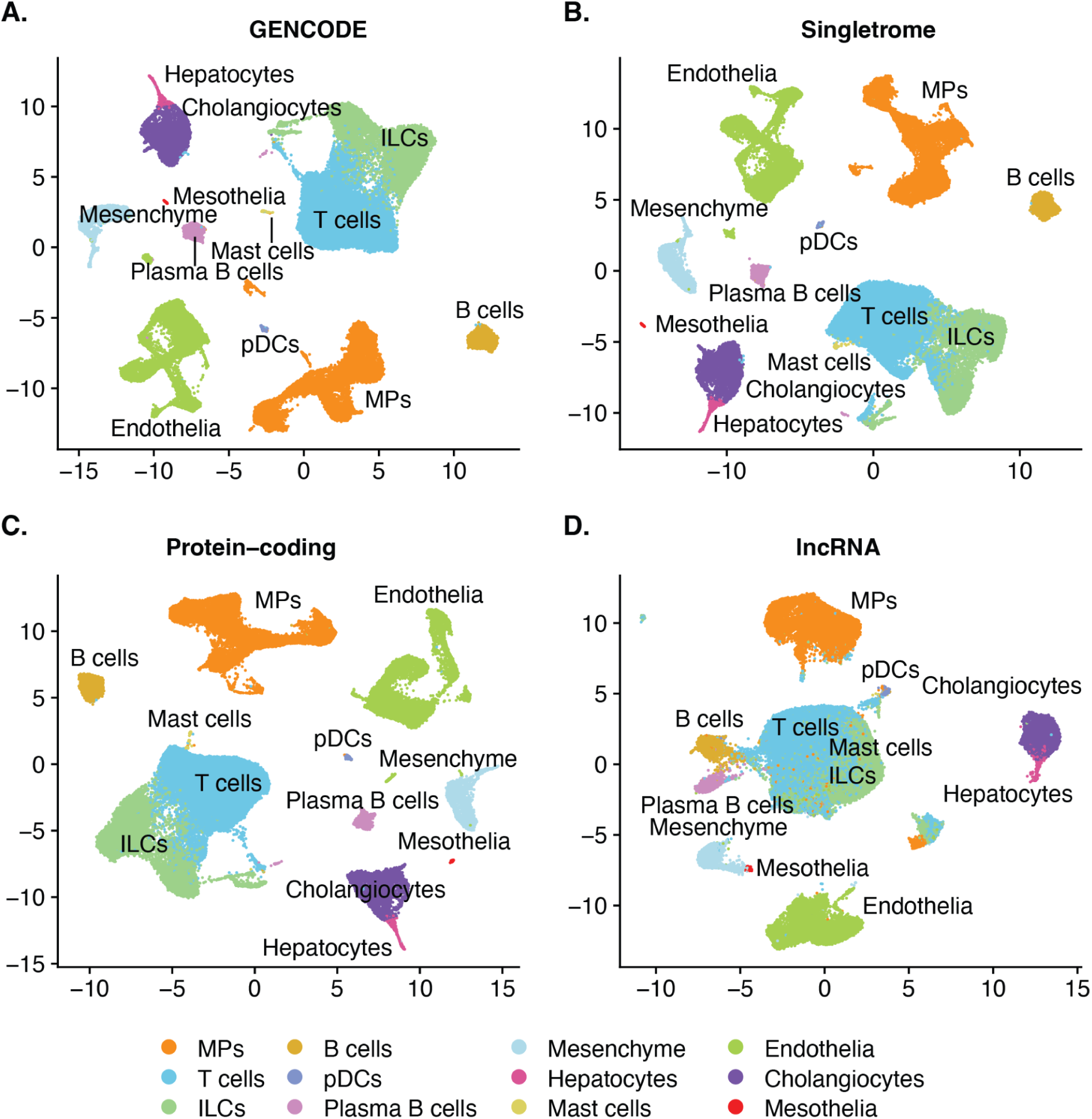
lncRNAs alone predict most clusters and cell types in single cell data of liver set 2. scRNA-seq data of liver set 2 from healthy and cirrhotic cells were mapped using annotation from **(A)** GENCODE, **(B)** Singletrome, **(C)** only protein-coding genes in Singletrome, and **(D)** only lncRNAs in Singletrome. The labels for each cell were retained from the original publication. For this analysis, Singletrome only contains lncRNAs that meet all filters developed with analysis of PBMCs and applied to data from liver set 2.

**Supplementary Figure 11.**
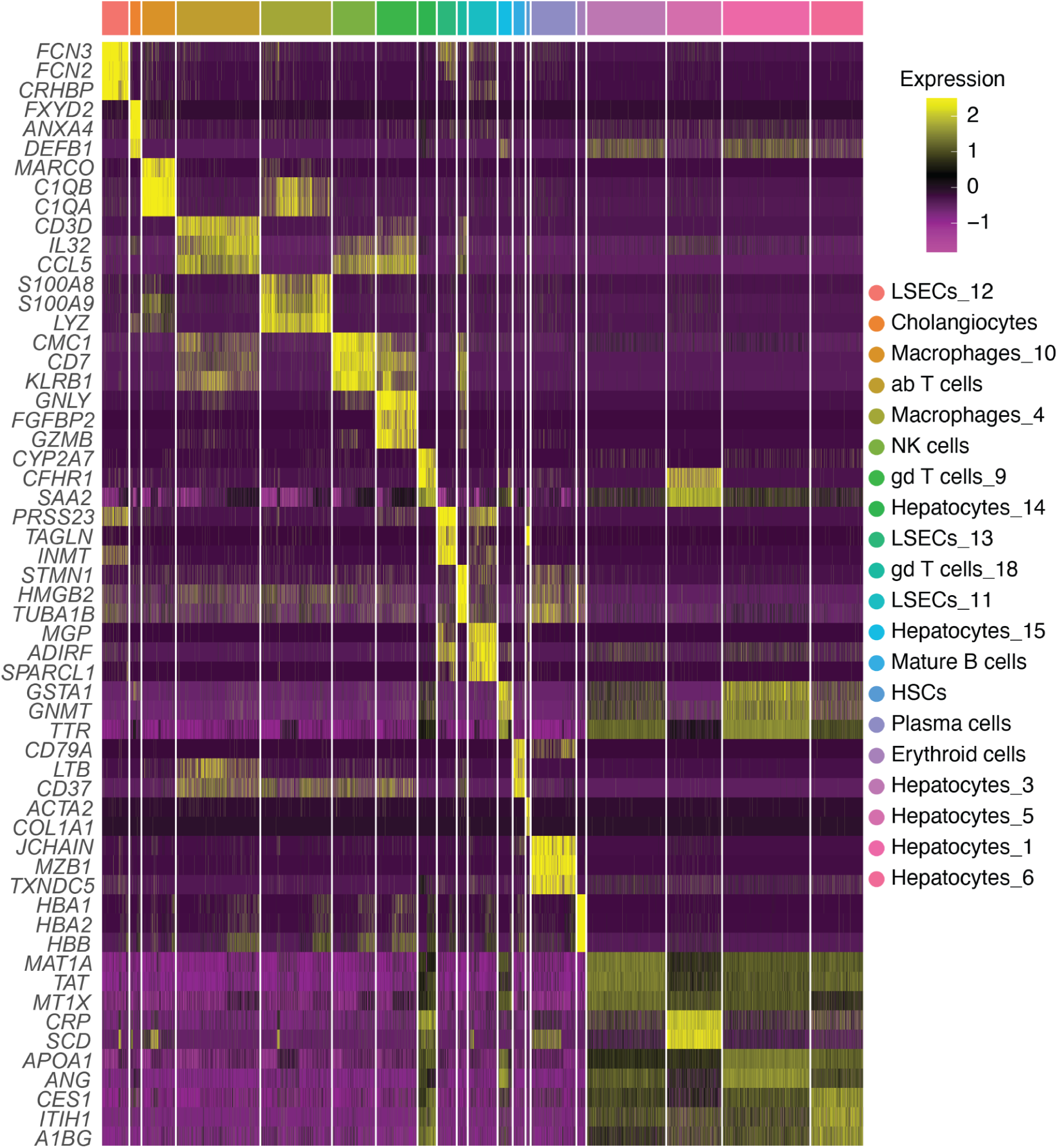
Protein-coding based cell type markers for liver set 1. The heatmap displays the top differentially expressed protein-coding genes (y-axis) for each cell type in liver set 1. Cell types are indicated by color at the bar above the heatmap, and the key is displayed to the right. Expression level is indicated by Z-score.

**Supplementary Figure 12.**
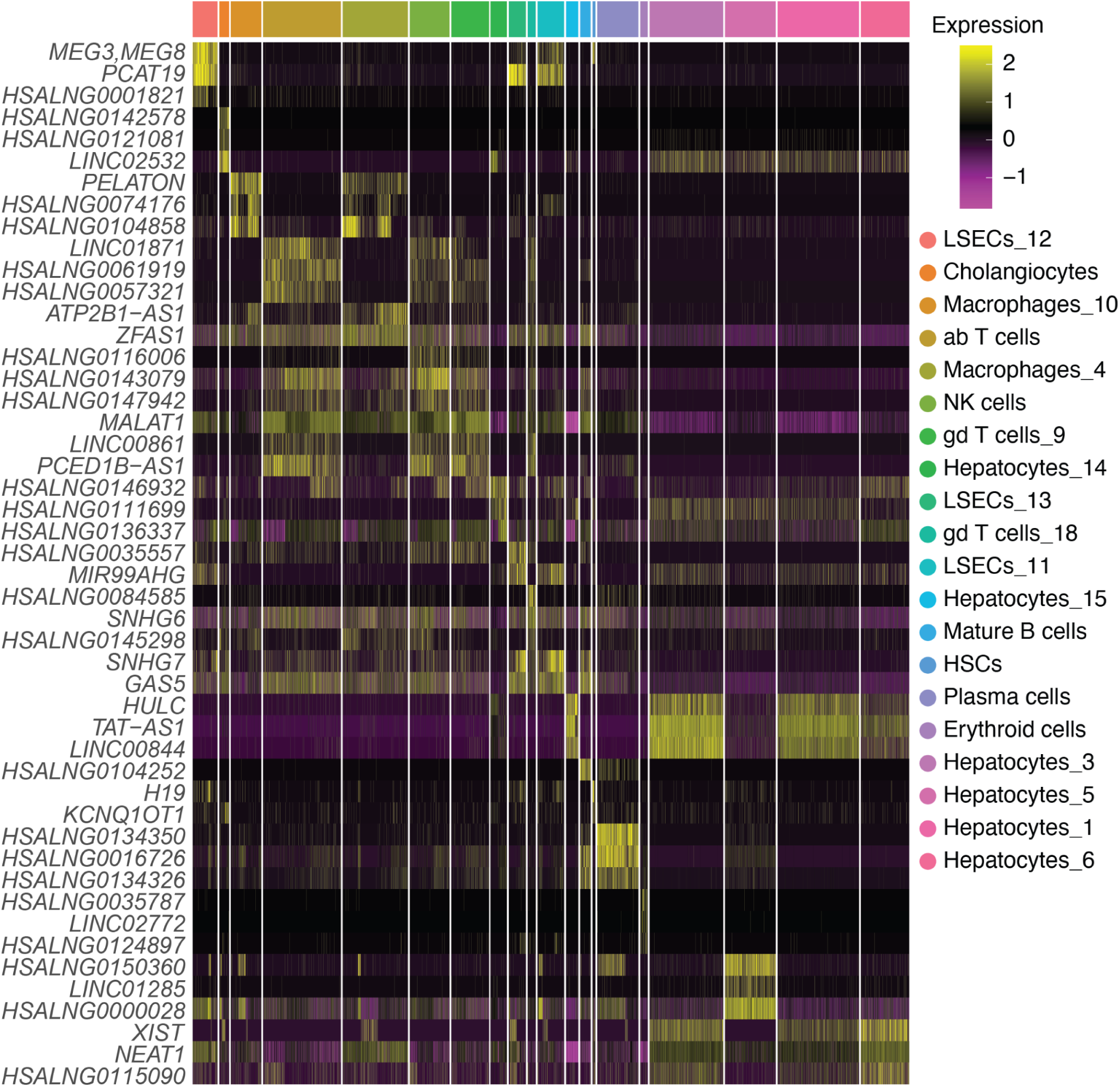
lncRNA based cell type markers for liver set 1. The heatmap displays the top differentially expressed lncRNA genes (y-axis) for each cell type in liver set 1. Cell types are indicated by color at the bar above the heatmap, and the key is displayed to the right. Expression level is indicated by Z-score.

**Supplementary Figure 13.**
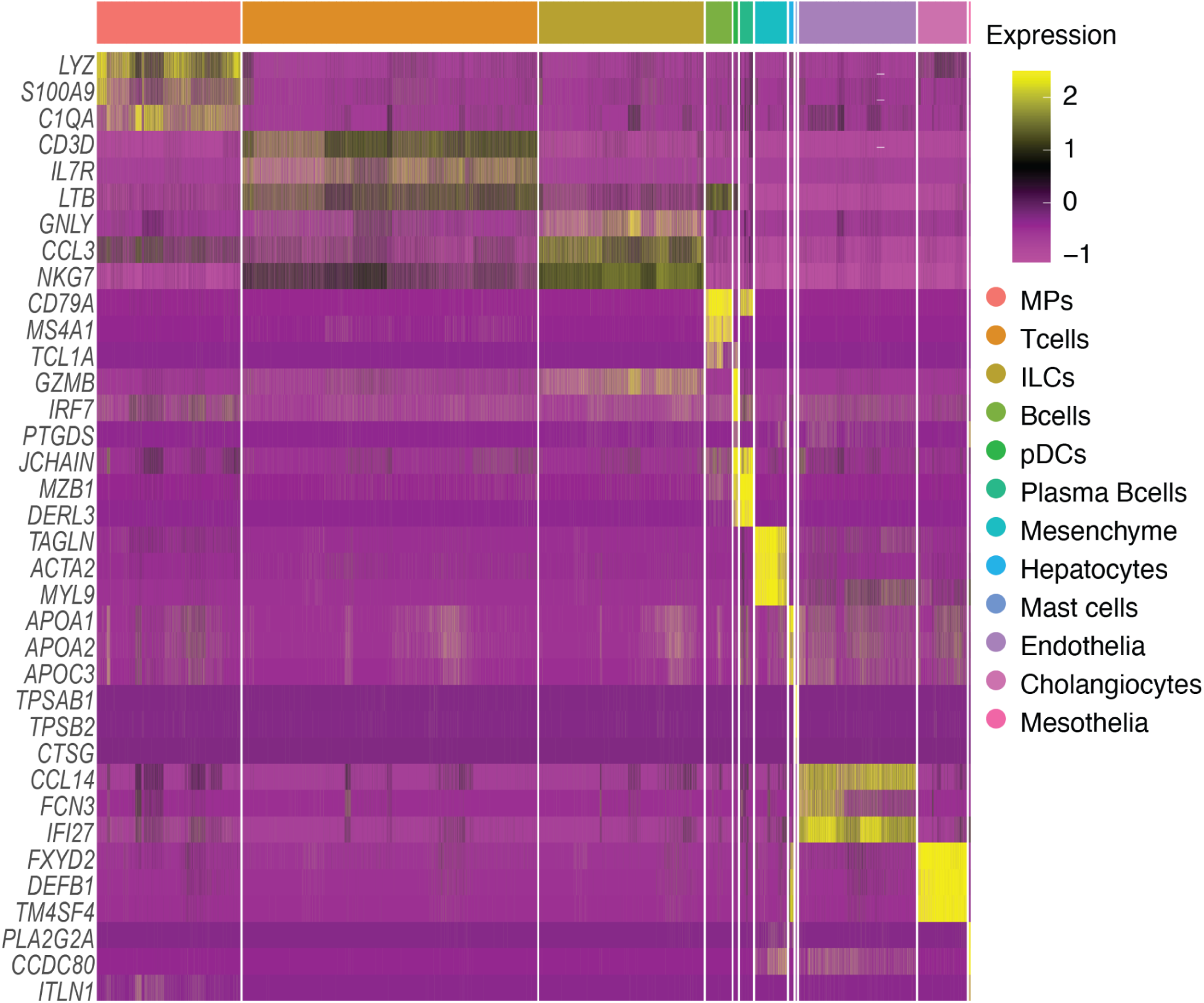
Protein-coding based cell type markers for liver set 2. The heatmap displays the top differentially expressed protein-coding genes (y-axis) for each cell type in liver set 2. Cell types are indicated by color at the bar above the heatmap, and the key is displayed to the right. Expression level is indicated by Z-score.

**Supplementary Figure 14.**
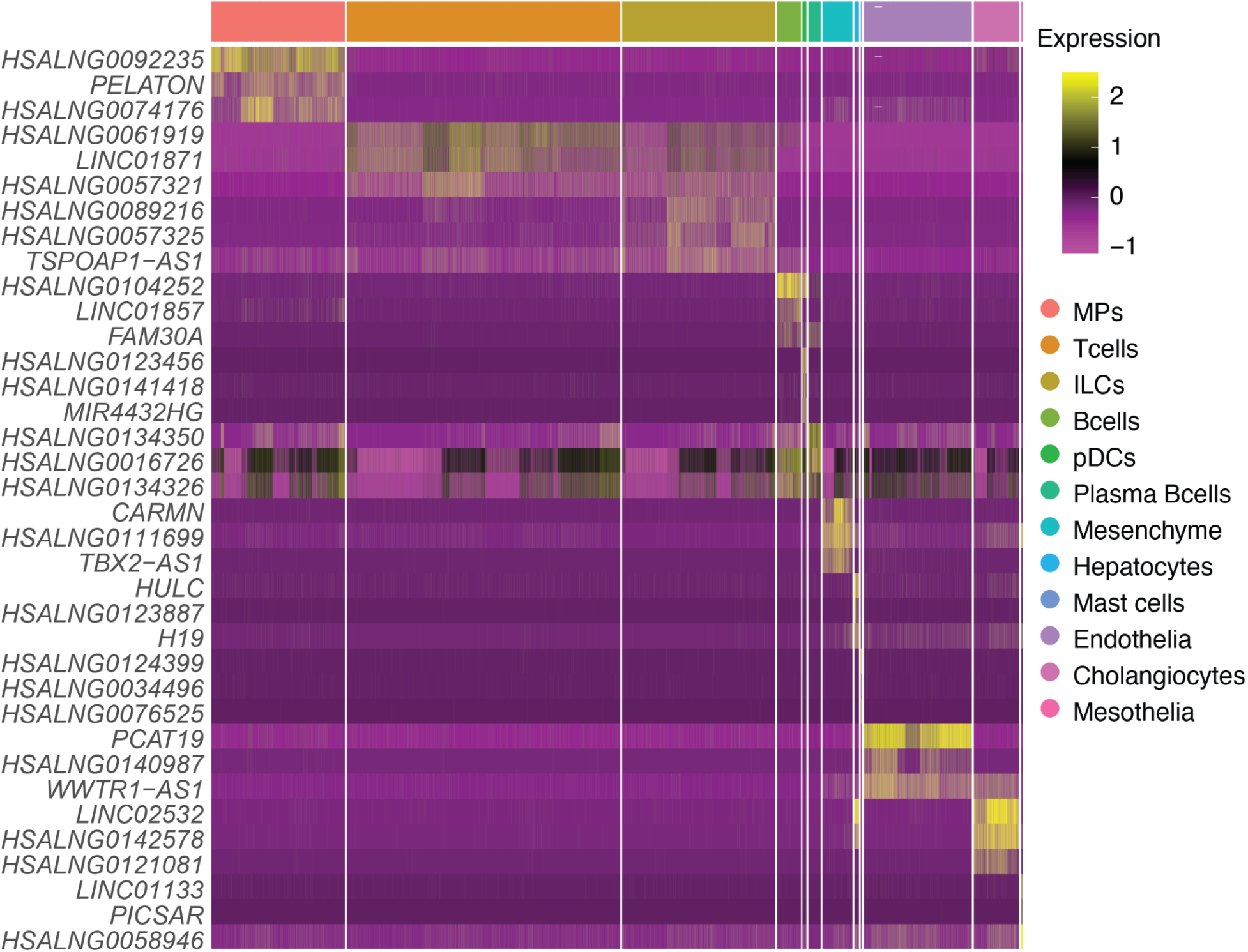
lncRNA based cell type markers for liver set 2. The heatmap displays the top differentially expressed lncRNA genes (y-axis) for each cell type in liver set 2. Cell types are indicated by color at the bar above the heatmap, and the key is displayed to the right. Expression level is indicated by Z-score.

**Supplementary Figure 15.**
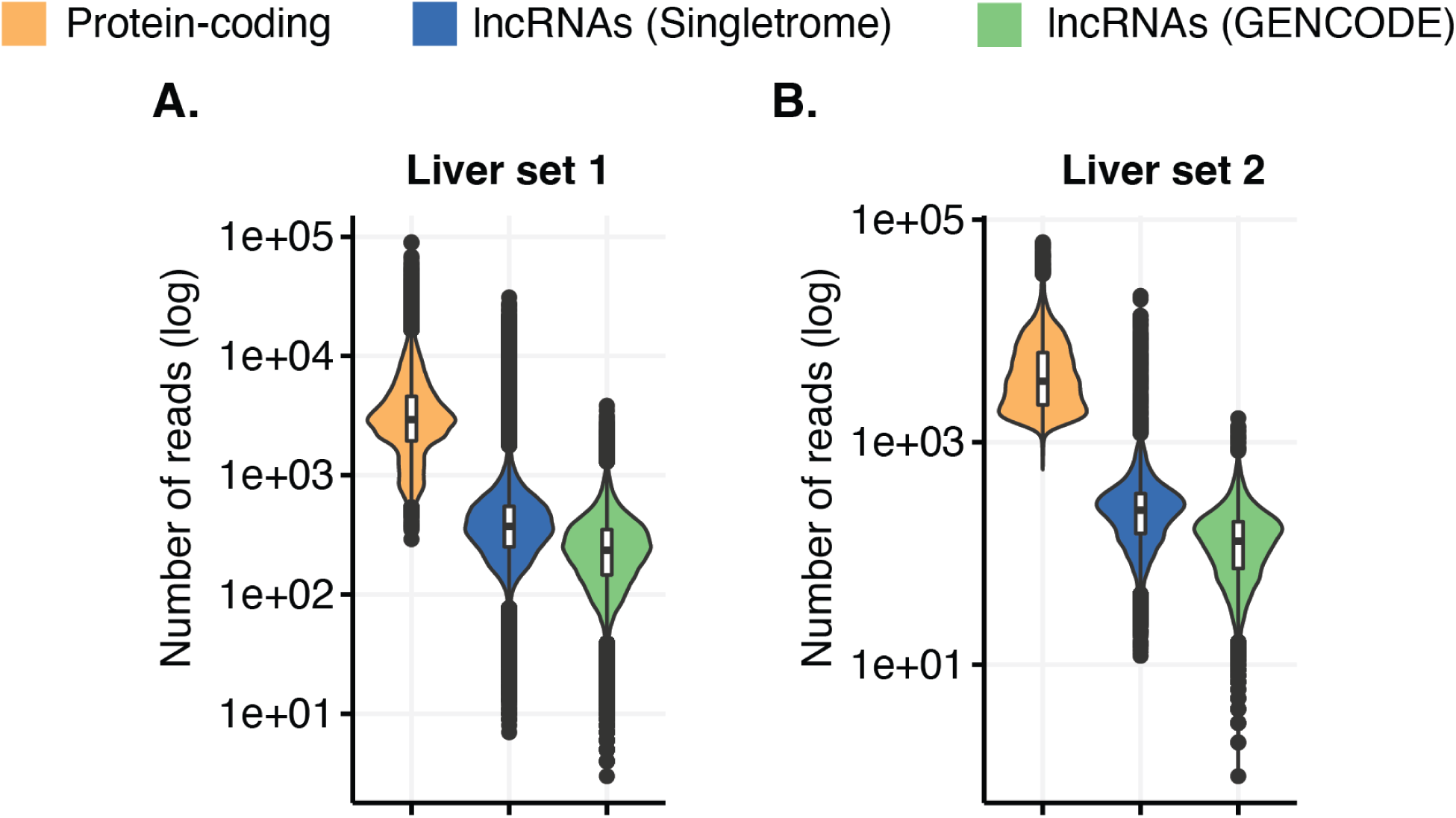
Expression of quality filtered lncRNAs compared to protein-coding genes in liver. The total number of mapped reads per cell (y-axis, log scale) is quantified for protein-coding genes (orange), lncRNA genes from Singletrome (blue), and lncRNA genes from GENCODE (green) in liver set 1 **(A)** and liver set 2 **(B).**

**Supplementary Figure 16.**
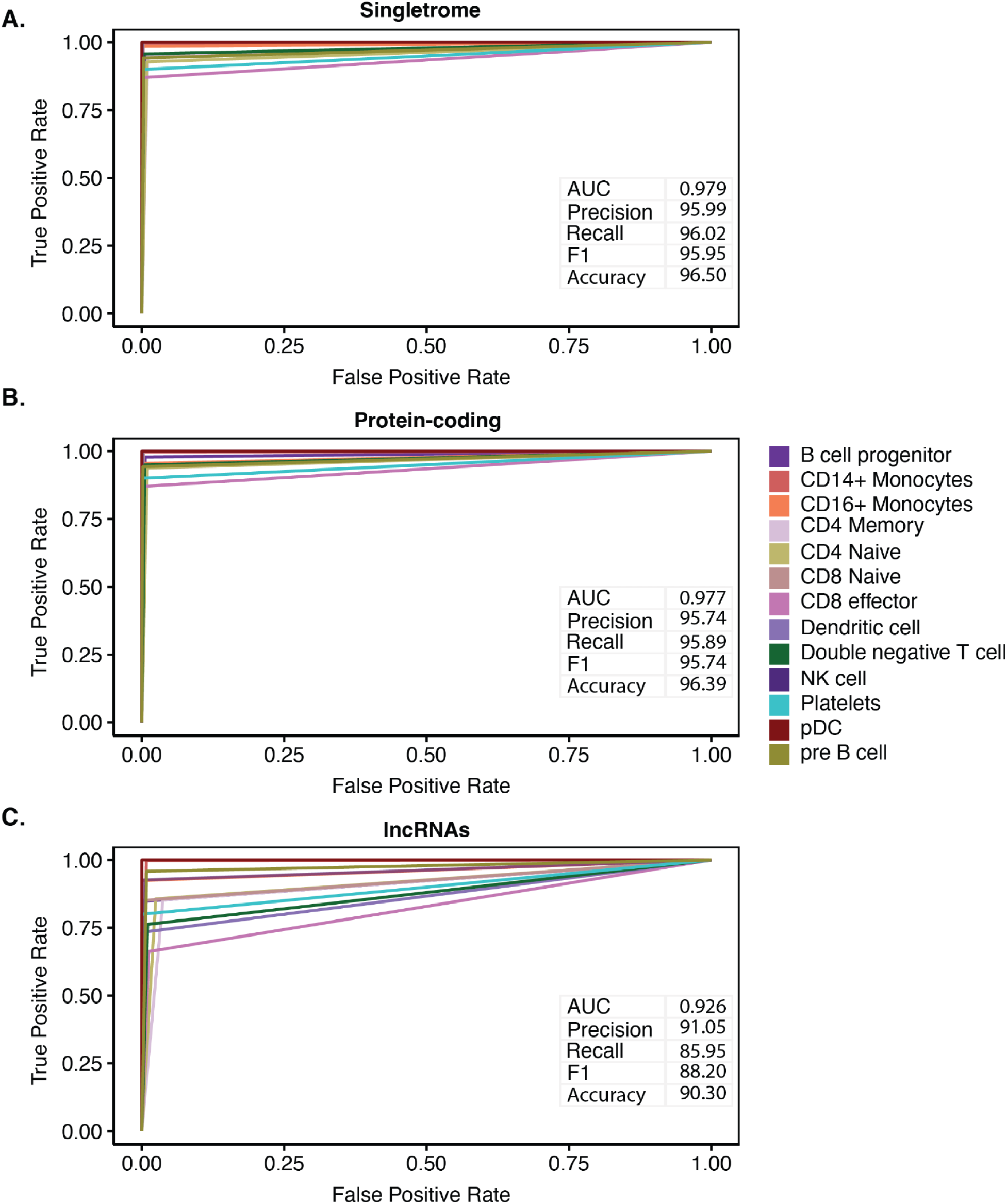
Cell type prediction for PBMCs. Receiver-operating characteristic (ROC) curve showing true and false positive rates for cell type prediction based on the expression of all genes in Singletrome **(A),** protein-coding genes alone **(B),** and lncRNA genes alone **(C)**. Cell types are indicated by color of the line, and the key is displayed to the right. The table inside the panel of each (A-C) shows the AUC, precision (%), recall (%), F1 (%), and accuracy (%) for cell type prediction of PBMCs.

**Supplementary Figure 17.**
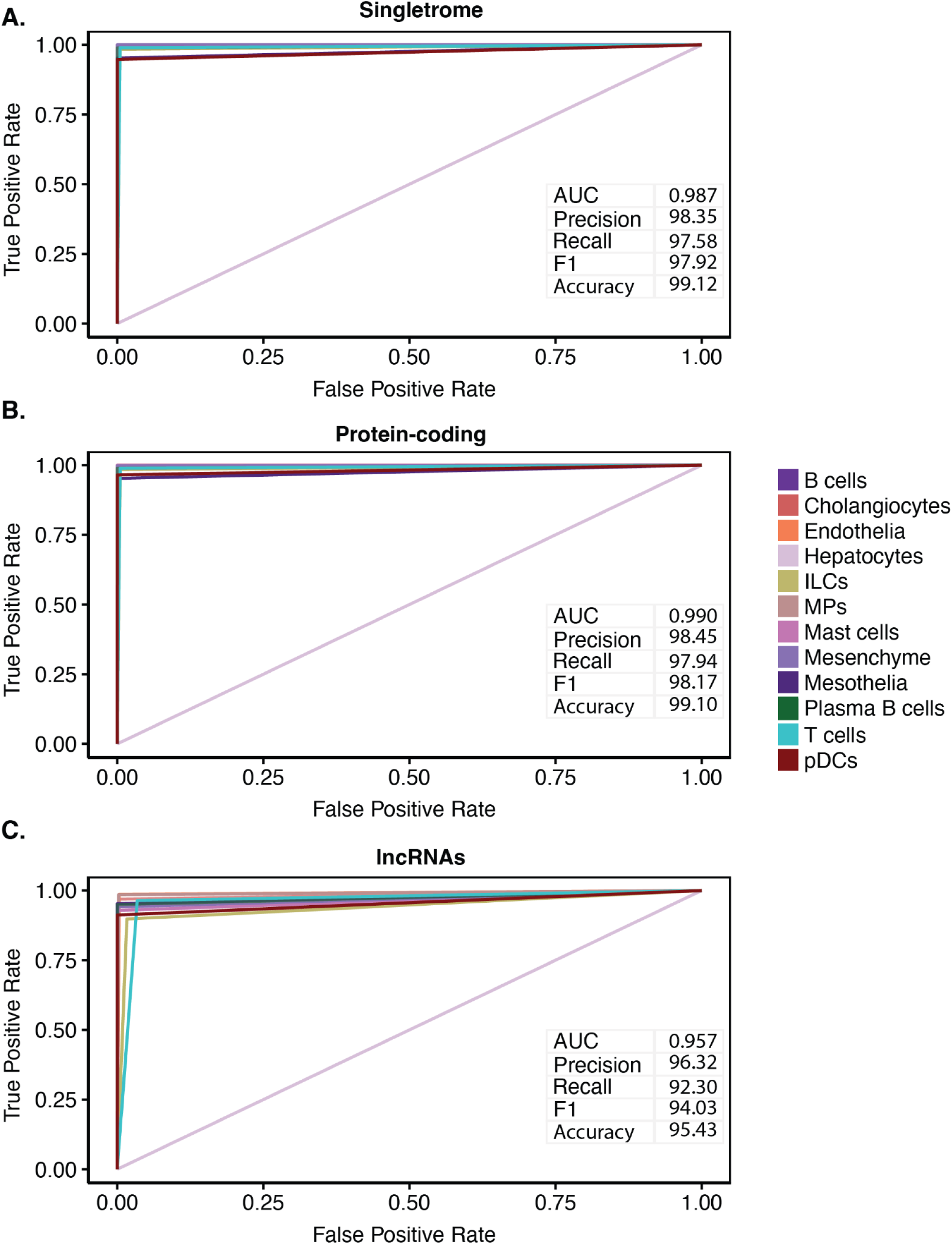
Cell type prediction for liver set 2. Receiver-operating characteristic (ROC) curve showing true and false positive rates for cell type prediction based on the expression of all genes in Singletrome **(A),** protein-coding genes alone **(B),** and lncRNA genes alone **(C)**. Cell types are indicated by color of the line, and the key is displayed to the right. The table inside the panel of each (A-C) shows the AUC, precision (%), recall (%), F1 (%), and accuracy (%) for cell type prediction of liver set 2.

**Supplementary Figure 18.**
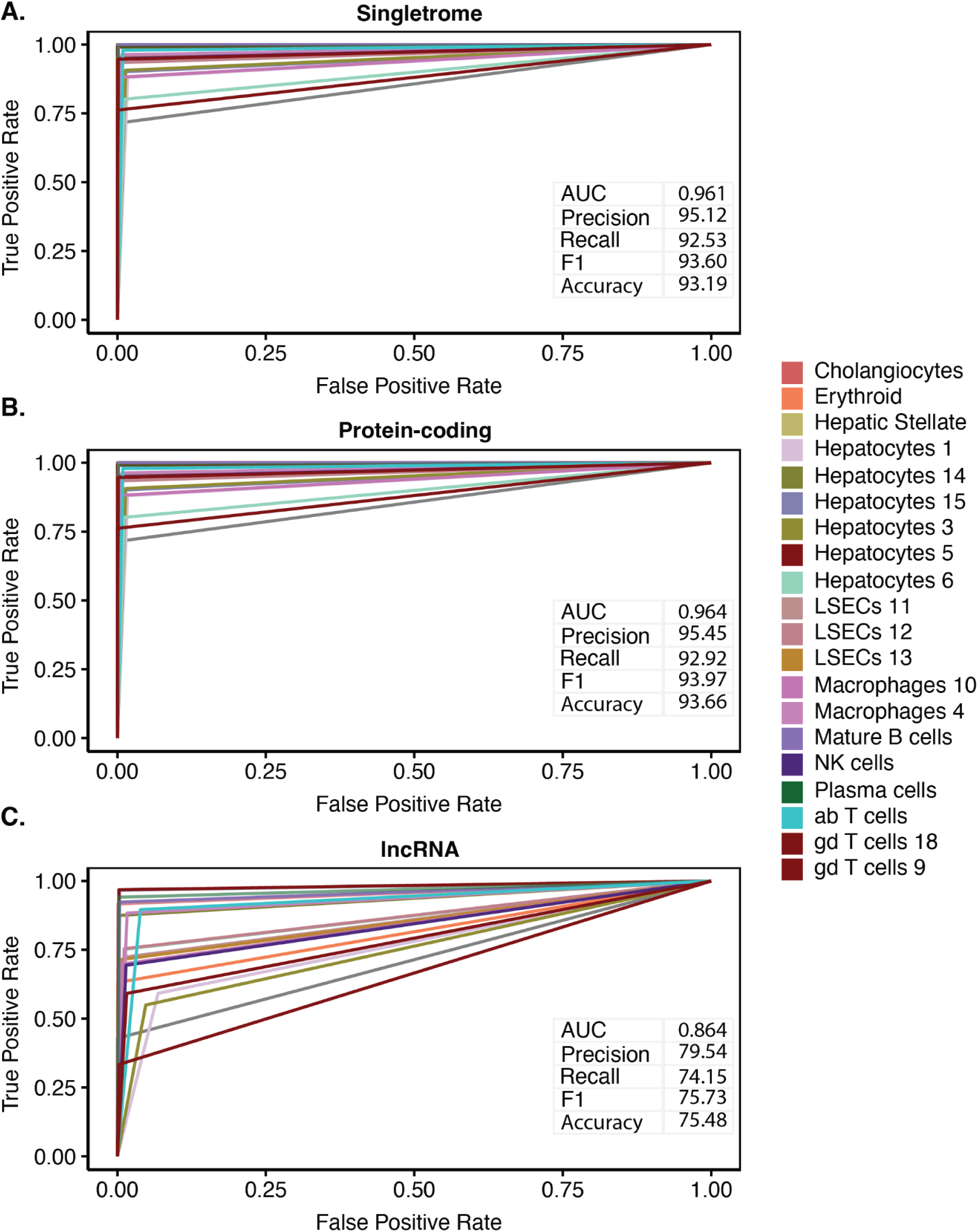
Cell type prediction for liver set 1. Receiver-operating characteristic (ROC) curve showing true and false positive rates for cell type prediction based on the expression of all genes in Singletrome **(A),** protein-coding genes alone **(B),** and lncRNA genes alone **(C)**. Cell types are indicated by color of the line, and the key is displayed to the right. The table inside the panel of each (A-C) shows the AUC, precision (%), recall (%), F1 (%), and accuracy (%) for cell type prediction of liver set 1.

**Supplementary Figure 19.**
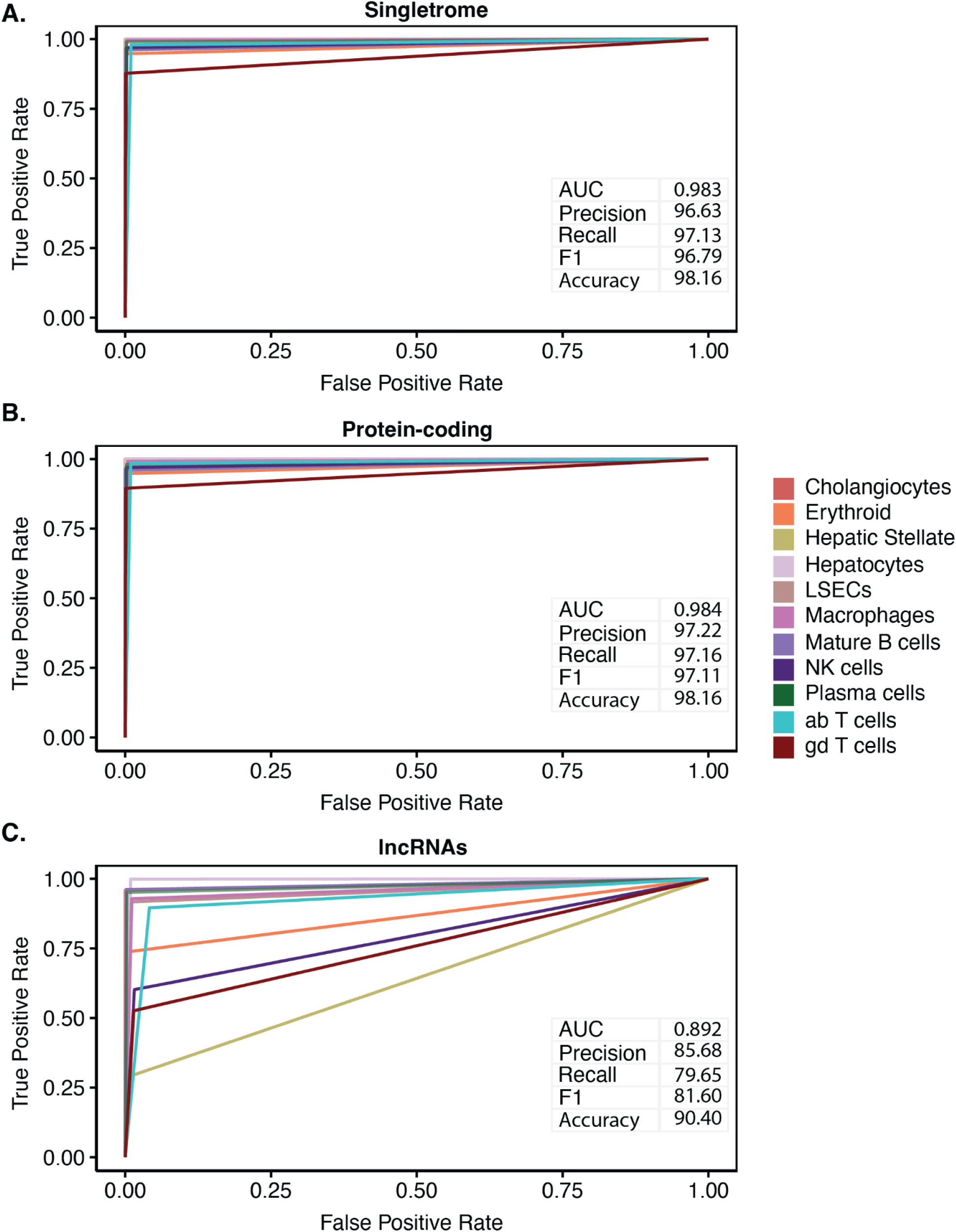
Cell type prediction for liver set 1 (sub-clusters merged to the same cell type). Receiver-operating characteristic (ROC) curve showing true and false positive rates for cell type prediction based on the expression of all genes in Singletrome **(A),** protein-coding genes alone **(B),** and lncRNA genes alone **(C)**. Cell types are indicated by color of the line, and the key is displayed to the right. The table inside the panel of each (A-C) shows the AUC, precision (%), recall (%), F1 (%), and accuracy (%) for cell type prediction of liver set 1.

**Supplementary Figure 20.**
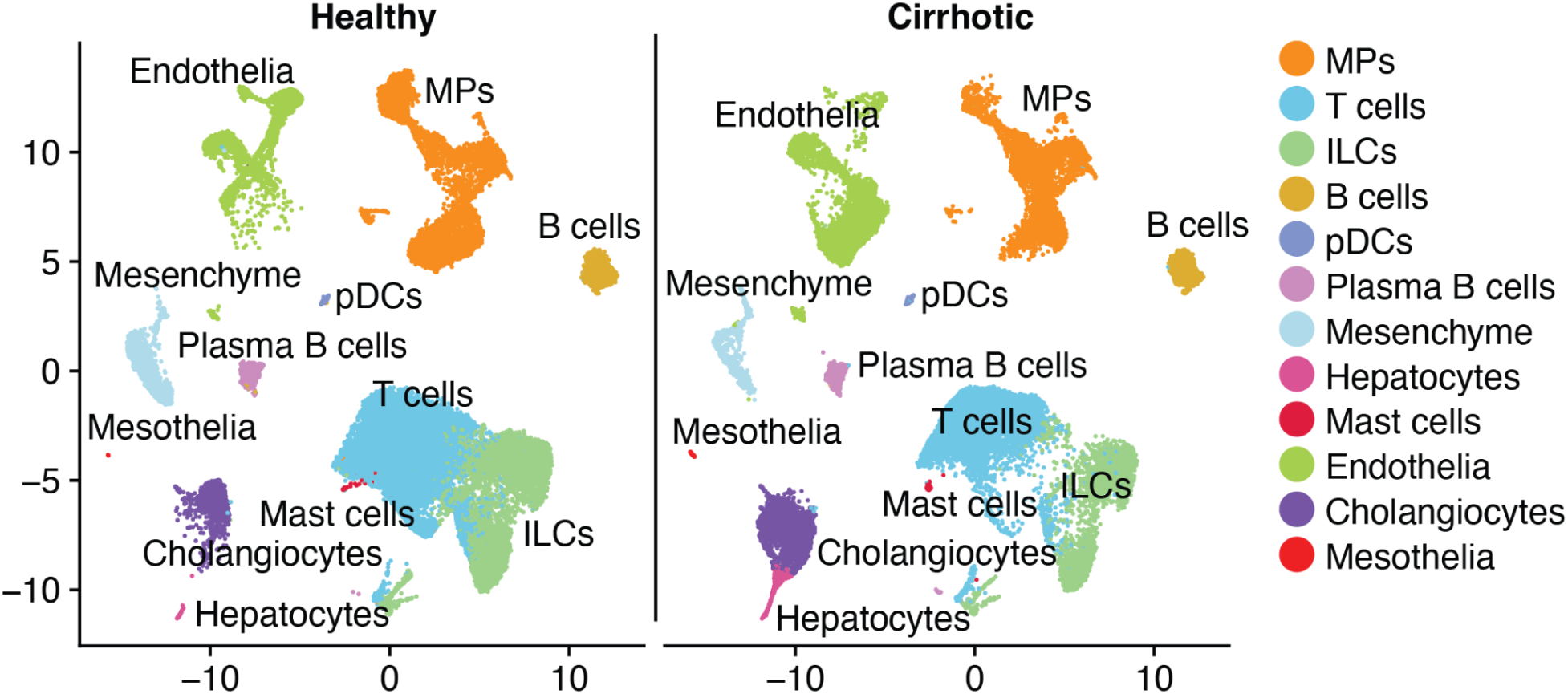
Singletrome cell type map in healthy and cirrhotic liver. scRNA-seq data of liver set 2 (GSE136103(Ramachandran et al. 2019)) were mapped using annotations from Singletrome. The labels for each cell were retained from the original publication. Cells were clustered based on all the genes in Singletrome and annotated by condition healthy (left) and cirrhotic liver (right). For this analysis, Singletrome contains all the protein-coding genes and only lncRNAs that meet all the described filters in the section ‘Quality control of lncRNA mapping’.

**Supplementary Figure 21.**
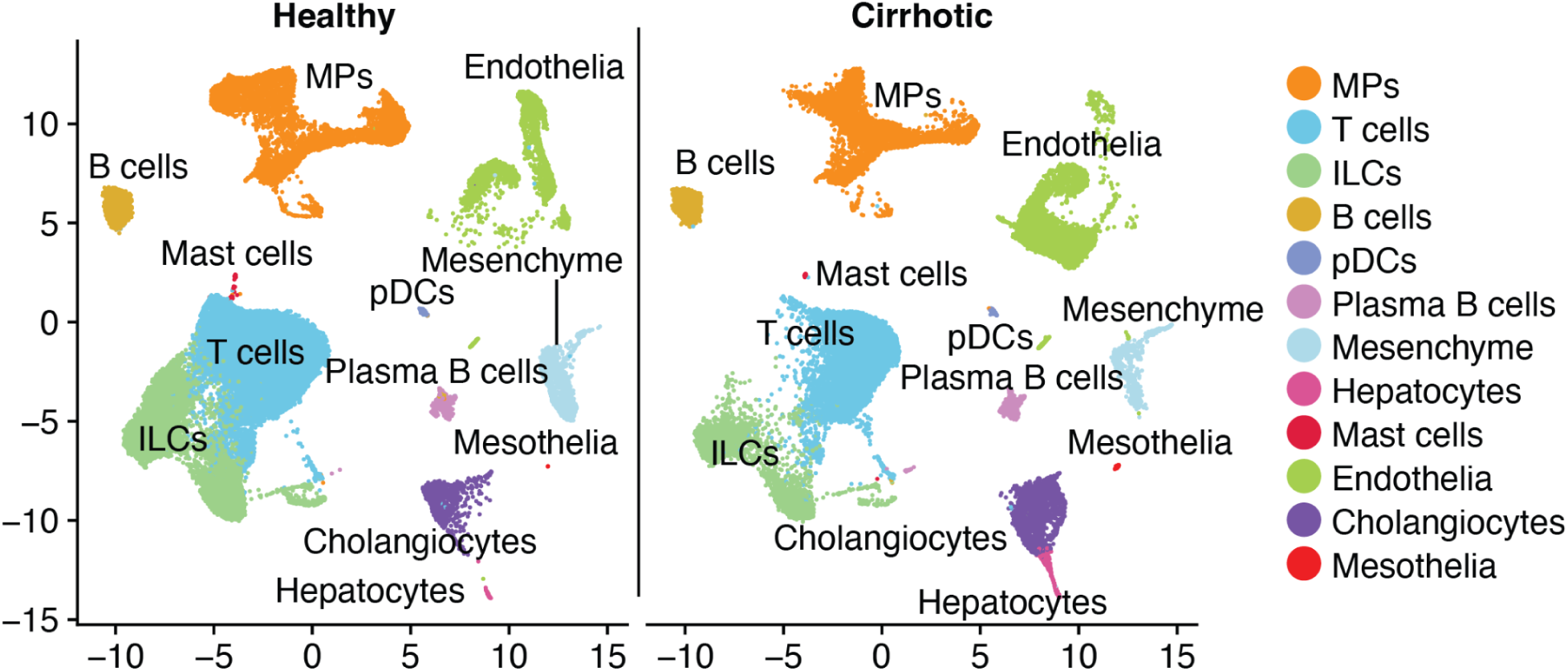
Protein-coding cell type map in healthy and cirrhotic liver. scRNA-seq data of liver set 2 (GSE136103(Ramachandran et al. 2019)) were mapped using annotations from Singletrome. The labels for each cell were retained from the original publication. Cells were clustered based on protein-coding genes from Singletrome and annotated by condition healthy (left) and cirrhotic liver (right).

